# Involvement of remote regions in sustained, but not transient, epileptic activities in the kainate mouse model of temporal lobe epilepsy

**DOI:** 10.1101/2023.09.14.557684

**Authors:** Guru Prasad Padmasola, Fabien Friscourt, Karl Schaller, Christoph M Michel, Laurent Sheybani, Charles Quairiaux

## Abstract

Animal and human studies have shown that the seizure-generating region is vastly dependent on distant neuronal hubs that can decrease duration and propagation of ongoing seizures. However, we still lack a comprehensive understanding of the impact of distant brain areas on specific interictal or ictal epileptic activities (e.g., isolated spikes, spike trains, seizures). Such knowledge is critically needed since all kinds of epileptic activities are not equivalent in terms of clinical expression and impact on the progression of the disease. We used surface, high-density EEG and multisite intracortical recordings, combined with pharmacological silencing of specific brain regions in the well-known kainate mouse model of temporal lobe epilepsy. We tested the impact of selective regional silencing on the generation of epileptic activities within a continuum ranging from very transient to more sustained and long-lasting discharges reminiscent of seizures. Silencing the contralateral hippocampus completely suppresses sustained ictal activities in the focus, as efficiently as silencing the focus itself, but while focus silencing abolishes all focal activities, contralateral silencing fails to control transient spikes. In parallel, we observed that sustained epileptic discharges in the focus are preceded by contralateral firing and more strongly phase locked to bi-hippocampal delta/theta oscillations than transient spiking activities, reinforcing the presumed dominant role of the contralateral hippocampus in promoting long-lasting, but not transient, epileptic activities. Altogether, our work provides suggestive evidence that the contralateral hippocampus is necessary for the interictal-to ictal-state transition and proposes that cross-talk between contralateral neuronal activity and ipsilateral delta/theta oscillation could be a candidate mechanism underlying the progression from short to long-lasting epileptic activities.

**Key Points:**

- We study how regions remote from the focus influence epileptic activities in the kainate mouse model of temporal lobe epilepsy.
- The contralateral hippocampus plays a decisive role in the initiation of sustained epileptic activities
- Integration of contralateral activities and bi-hippocampal delta/theta oscillations precedes focal paroxysmal activities
- We propose that a large-scale epileptic network might be necessary for the transition from interictal to ictal states

## Introduction

Focal epilepsies are believed to rely on the activity of an epileptic focus where seizures originate. However, recent works on the impact of remote regions on the activity of the focus ^1–4^ have raised the hypothesis that the focus should rather be considered a “focus-network”, implying that several interconnected key regions are involved in and can modulate the generation of epileptic activities ^5–9^. For example, it has been shown that remote regions can contribute to decrease the duration of epileptic seizures ^2,3^. Still, the network mechanisms that favor the generation of the different types of epileptic activities and in particular the expression of transient vs sustained epileptic activities remain in general vastly unknown. This is crucial, since the impact of interictal epileptiform discharges (IEDs), at one extreme of the spectrum, is very different from the clinical semiology of seizures. A more comprehensive understanding of the interactions between nodes of a large-scale epileptic network is also necessary to capture the complexity of seizure dynamics ^9,10^, the networks distribution of IEDs as well as certain cases of post-surgery seizure relapses ^11^.

We thus aim to investigate the network aspect of ictogenic mechanisms and, more specifically, address the open question of whether the transition from interictal states, characterized by transient epileptic activities, to sustained epileptic activities, depends on brain areas remote from the primary focus. In the well-known kainate mouse model of unilateral temporal lobe epilepsy ^4,12–14^, we used surface, high-density EEG (HD-EEG) and bi-hippocampal intracortical EEG (iEEG) to reveal a spectrum of epileptic activities, from short, isolated spikes to long-lasting hippocampal paroxysmal discharges. Using selective pharmacological silencing, we studied the impact of different target brain regions (ipsilateral hippocampus and frontal motor cortex and contralateral hippocampus), on each of these classes of epileptic activities. Finally, we analyzed bi-hippocampal coupling (including multi-unit activity) with the aim of understanding the mechanisms underlying the association between contralateral hippocampal silencing and abortion of interictal to ictal states transition.

Our work establishes that escalation towards long-lasting, sustained discharges reminiscent of ictal activity is dictated by a fine balance between both hippocampi, with a particular role of the contralateral hippocampus in long-lasting discharges.

## Methods

Forty-seven male C57BL/6j adult mice (10-11 weeks) were included. Experiments were performed in accordance with the Geneva and Switzerland animal care committee’s regulations. Part of the intracortical data comes from experiments previously published ^4,15,16^.

### Surgeries

Head-fix surgeries were performed as previously described ^4,15^ (further detailed in Supporting Information). Injectable anesthesia was used during the surgery (Medetomidine: 0.5 mg/kg, Midazolam: 5 mg/kg, Fentanyl: 0.05 mg/kg, i.p.) and for awakening (Flumazenil: 0.5 mg/kg, Atipamezol: 2.5 mg/kg, Naloxone: 1.2 mg/kg, subcutaneous). Animals were placed back in their cage for 7 days before kainate injections. Antibiotics (trimethoprime-sulfamethoxazole) and anti-inflammatory analgesics (Ibuprofen, Paracetamol) were provided in drinking water during first 48h.

### Kainate and tetrodotoxin (TTX) injections

Kainate (70 nl, 5 mM in NaCl 0.9%) was injected under isoflurane anaesthesia with a pulled glass capillary in the left dorsal hippocampus (Mediolateral 1.6 mm, Anteroposterior 1.8 mm, Depth 1.9 mm), inducing a status epilepticus. TTX (0.15 µl; 0.3 mM in NaCl 0.9%) injections were made to decrease neuronal activities in 3 regions that express epileptiform discharges ^4,15,16^: focus (AP −2.67 mm, ML 2.5 mm, depth 1.72 mm and 1.22 mm, angle 20°), contralateral hippocampus (HD-EEG: AP 2.54 mm, ML 2.10 mm, depth 2.0; iEEG: AP −2.67 mm, ML −2.5 mm, depth 1.72 mm and 1.22 mm, angle 20°) or ipsilateral frontal cortex (AP 1.98 mm, ML 1.3 mm, depth 0.5 mm). We previously showed that such TTX injections had localized effects on neuronal activities ^4^.

### Electrophysiological recordings

Recordings were acquired (Digital LynxSX, Neuralynx) after 3-4 days of training to the head-fixed apparatus. HD-EEG was recorded (at 4 KHz, low pass 2 KHz) with stainless electrodes (Fig.S1) as previously described ^4,17^ and re-referenced to the average reference offline. iEEG was recorded (at 16 KHz, low pass 8 KHz) with longitudinal 16-electrode probes (Neuronexus) implanted in both hippocampi (AP, ML, depth, angles: 2.67, 2.5, 2.02, 20°; Fig.S2), against cerebellum electrodes. Animals were briefly anesthetized with isoflurane to allow electrode positioning; recordings were performed after complete awakening.

### Classification of epileptiform discharges

We applied semi-automated procedures combining algorithm-based fast-ripples (FRs) detection and visually detected epileptiform discharges as previously described ^4,16^. Overriding FRs almost systematically accompany epileptiform discharges, occurring during the spike component, and vice-versa ^4,18,19^. The automated FRs detection algorithm allows to standardize detection and eliminate examiner bias, although it is possible that some epileptiform discharges not combined with FRs are discarded.

<u>Focus epileptiform discharges</u> were classified using the consensual parameters of duration and intrinsic spiking frequencies. Classifications vary across studies but a consensus emerges to consider longer lasting focus bursts with high spiking rates, accompanied or not by behavioral immobility and muscular twitches, as ictal-like patterns reminiscent of focal seizures, although different terms have been used in the literature ^4,12,14,20–22^. Shorter events are commonly considered as IEDs. Generalized tonico-clonic seizures are also present in the model ^4,12,23^, however more rarely detected in our 1h recordings.

For HD-EEG, a marker is positioned on the positive peak of the spike waveform associated with each FR detected above the focus (Fig.S1). For iEEG, the marker is positioned at the peak of the spike waveform in the dentate gyrus (DG, Fig.S2). To avoid counting twice volume-conducted FRs, each FR is counted only once even when detected simultaneously on several channels. Peak-to-peak inter-spike intervals are then calculated in order to classify epileptic patterns as follows. Discharges that are neither preceded nor followed by another spike for at least 1 sec are classified as <u>isolated spikes</u> (IS). Events with more than two successive spikes within a second are further separated according to spike rates and event duration. Events that never display >5 spikes/second are named <u>spike trains</u> (STs); the first detected spike not followed by another spike for at least one second marks their termination. Events that reach >5 spikes/second but do not exceed 10 seconds are classified as <u>short</u> <u>hippocampal paroxysmal discharges</u> (sHPDs), those that exceed 10 seconds are classified as <u>ictal hippocampal paroxysmal discharges</u> (iHPDs). This type of long electrographic paroxysmal event is reminiscent of focal seizures, although concomitant epileptiform discharges can be occasionally observed in the contralateral hippocampus.

<u>Network IEDs</u> were identified visually and quantified as separate events. Network IEDs correspond to the previously described generalized spikes in ^4^, i.e., IEDs invading a large-scale network and characterized by two spiking bursts separated by flattening of the EEG of approximately 250 ms.

### Signal analysis

<u>Frequency power</u> was calculated using ft_freqanalysis in fieldtrip ^24^. For background power, 50 epochs were selected per recording such that they were 2 sec away from any detected epileptic events. Power was calculated using Morlet wavelet analyses for time-frequency plots (Fig.1, Fig.S1).

**Figure 1.**
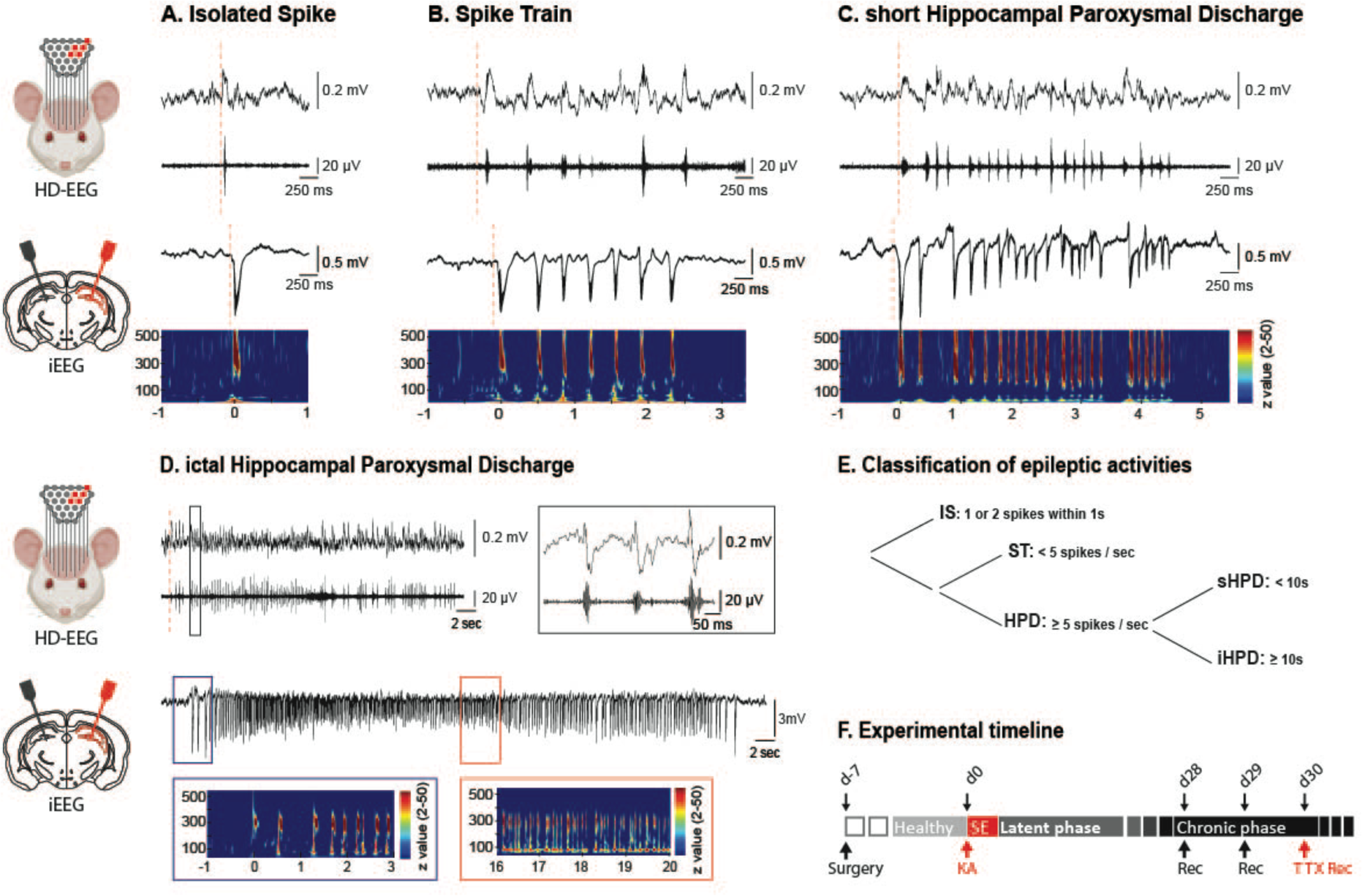
Classification of epileptiform discharges recorded from the epileptic focus based on duration and spike rates during surface HD-EEG and bi-hippocampal iEEG recordings. (A-D) Illustrative examples of the 4 types of focal epileptiform discharges detected during one HD-EEG recording session (upper panels, all events from electrode e19, see supplementary fig. S1 for the electrode map) and during one iEEG recording session (lower panels, electrode in the DG of the kainate injected hippocampus). Left cartoons illustrate the recording methods; red channels are above (HD-EEG) or inside (iEEG) the epileptic hippocampus. In HD-EEG panels, the *top trace shows the raw LFP and the bottom trace the filtered LFP (250-500 Hz), which reveals the FRs overriding the spikes. FRs are readily visible in the raw LFP signal, as illustrated by the zoom in the black rectangle of panel D (See also supplementary Fig. S1). In iEEG panels, raw traces and* corresponding time-frequency plots (0-500 Hz; time axis in sec; z-scored against background; z-scores with values < 2 are set to 0) are shown, revealing that the sharp discharges are characterized by increased power in both the epileptic spike (15-30 Hz) and the fast-ripple band (250-500 Hz). Observe the change in frequency of spikes between spike trains and paroxysmal discharges (sHPD and iHPD). The onset of focal iHPD (blue rectangle, panel D) is characterized by high amplitude low-frequency spiking activities which gradually change to lower amplitude higher-frequency spiking (red rectangle), reminiscent of an epileptic seizure. *Red dotted lines mark the onset of each event*. E. Classification rules of detected epileptic activities (see Methods) using the spike rates and event duration criterion. F. General experimental timeline. The surgery for the placement of the head-holder was made 7 days before the kainate injection. HD-EEG (n=31 mice) or iEGG (n=16 mice) were made during the chronic phase before and during TTX treatments.

<u>Multi-unit activity</u> (MUA) was detected in the iEEG recordings (600-6000 Hz) based on a threshold method ^15^.

<u>Phase-locking and phase-synchronization</u> analyses were performed in windows of 1 sec either preceding the statistical onsets of epileptiform discharges or in background periods. Data were first re-referenced to the cortical electrode of the probe and filtered in delta (0.5-4 Hz) and theta (4-12 Hz) frequency bands using a non-causal filter. Instantaneous amplitude and phase are obtained by Hilbert-transform. The statistical onset of an epileptic discharge was calculated based on a threshold detection method as in ^4,15^. The phase-locking of the epileptiform discharges was then obtained through intertrial coherence (ITC), which is defined as:

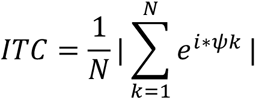

where N is the number of windows corresponding to epileptiform discharges or background periods, k the current window, and ψ the phase of the signal. ITC was calculated separately for each event and averaged across similar event type to obtain one ITC value per animal and event type. Channels with maximal theta power in the pyramidal region were selected for statistical analysis. Background periods of 1 second were randomly selected such that they were 2 seconds away from any epileptiform discharges and matched the number of epileptic discharges in each recording.

### Statistics

Statistical analyses were performed using Prism (GraphPad Software, Inc.). The normality of distributions was evaluated with the D’Agostino-Pearson method before adapted statistical tests were applied: t-test, Mann–Whitney (non-parametric distribution), Wilcoxon (non-parametric distribution paired groups), ANOVA (>2 groups; post hoc: Tukey’s multiple-comparison test), Kruskal–Wallis (>2 groups, nonparametric distribution unpaired groups; post hoc: Dunn’s multiple-comparison test). Bonferroni corrections were applied for multiple comparisons.

## Results

The kainate model reproduces a focal epileptic disease with different types of epileptiform discharges comprising long-lasting HPDs characterized by high-load spike bursts and reminiscent of electrographic focal seizures; these HPDs can be accompanied by behavioral immobility and spasms and occasionally generalize ^4,13,22,23^. In line with previous studies, we defined epileptiform discharges by the occurrence of epileptic spikes, i.e., sharp discharges peaking at 10-40 Hz, at electrodes positioned above or in the injected hippocampus and are classified by their duration and intrinsic spiking rates in 4 different types (Fig.1): isolated spikes (IS) and spike trains (ST), denoted together interictal epileptiform discharges, and short hippocampal paroxysmal discharges (sHPD) and ictal-HPD (iHPD) (see Methods). None of these events were detected in HD-EEG recordings made before kainate injections (d0) but they were all present in almost all kainate-injected animals recorded during the chronic stage at d28 or d29; only a few animals did not display sHPDs and iHPDs (5/31) but still exhibited IS or ST. The incidence of these epileptiform discharges was stable across recordings made at d28 and d29 (Fig.2). IS and ST were highly frequent (3-4/min and 0.4-0.5/min; see figures legends for detailed quantifications). About 6 ictal-HPDs were detected per hour, with a stable duration across recordings (median: d28=26.69 sec; d29=21.01 sec; p=0.08, paired t-test).

**Figure 2.**
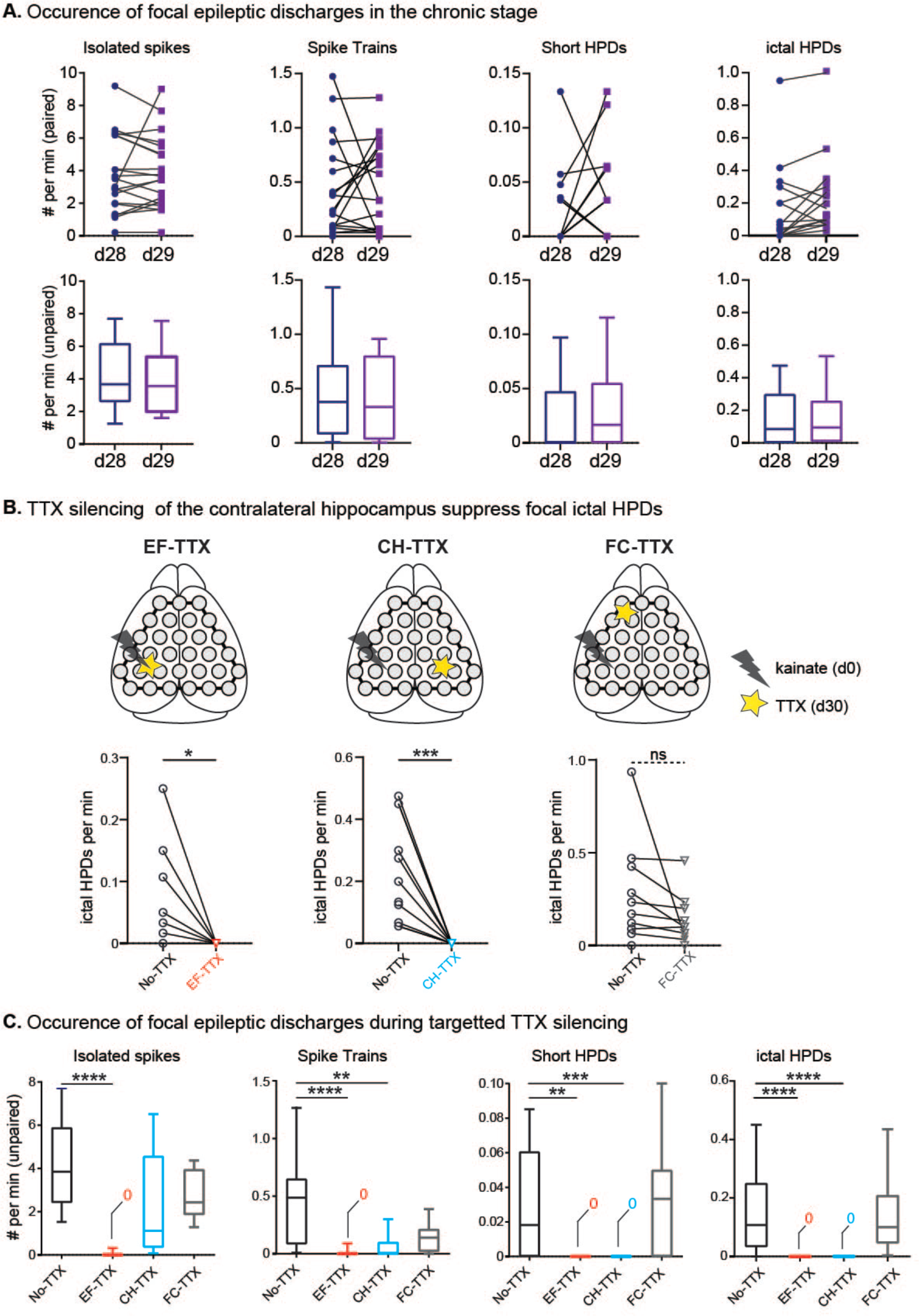
Effects of targeted silencing on the occurrence of focal epileptic activities during HD-EEG recordings. EF= epileptic focus; CH= contralateral hippocampus, FC= frontal cortex. A) Focal epileptic activities remained stable between recordings made on 2 successive days in the chronic stage (No-TTX condition). Above, paired comparisons of the number of epileptiform discharges per min in animals recorded at both d28 and d29 (n=20): isolated spikes (median and interquartile; D28: 3.25,1.96,6.18; D29: 3.56,1.97,5.37, p = 0.77, Wilcoxon test), spike trains (D28: 0.22,0.07,0.68; D292: 0.33,0.03,0.80, p = 0.64, Wilcoxon test), short HPDs (D28: 0, 0, 0.03; D29: 0.01, 0, 0.05, p = 0.26, Wilcoxon test) and ictal HPDs (D28: 0.01, 0, 0.2; D29: 0.09, 0, 0.25, p = 0.07, Wilcoxon test). Below, median number of epileptic activities per min across all animals (n=31 at d28, n=20 at d29) with q25-q75 IQ ranges (box plot) and 10-90 percentiles (whiskers): isolated spikes (Median and IQ; D28: 3.66,2.59,6.18; D29: 3.56,1.97,5.37, p = 0.63, Mann-Whitney test), spike trains (D28: 0.37,0.08,0.71; D29: 0.33,0.03,0.80, p = 0.68, Mann-Whitney test), short HPDs (D28: 0, 0, 0.04; D29: 0.01, 0, 0.05, p = 0.89, Mann-Whitney test) and ictal HPD (D28: 0.08, 0, 0.3; D29: 0.09, 0, 0.25, p = 0.96, Mann-Whitney test). B) In the same epileptic mice that were recorded at d28 or d29, no focal ictal HPDs were detected at d30 when TTX was injected in the epileptic focus (EF; n=10, median and IQ; No-TTX: 0.025,0,0.26; TTX: 0, p = 0.03, Wilcoxon test) or in the contralateral hippocampus (CH; n=11, No-TTX: 0.2,0.06,0.45; TTX: 0,p = 0.001, Wilcoxon test). The number of ictal HPDs recorded during TTX injection in the ipsilateral frontal cortex (FC) was not significantly different as compared to the recordings made without TTX (n=10, No-TTX: 0.2,0.08,0.43; TTX: 0.1,0.04,0.2, p = 0.13, Wilcoxon test). C) Median, IQ range and 10-90^th^ percentile of the occurrence of the 4 types of focal EAs across all groups (No-TTX=31 mice). As expected, TTX injection in the EF completely abolished all focal epileptic activities. Kruskal–Wallis with Dunn’s multiple-comparison tests confirm that TTX injections in the CH, but not in the FC, abolished ictal HPDs (No-TTX: 0.11,0.03,0.25; EF-TTX:0, p<0.0001; CH-TTX:0, p<0.0001; FC-TTX: 0.10,0.04,0.20, p>0.99;). Furthermore, these analyses revealed that short HPDs were also completely and specifically suppressed by CH silencing (No-TTX: 0.01,0,0.06; EF-TTX:0, p=0.0012; CH-TTX:0,p=0.0007; FC-TTX:0.03,0,0.05, p>0.99) while spike trains were significantly decreased but still present (No-TTX: 0.48,0.08,0.64; EF-TTX:0, p<0.0001; CH-TTX:0,0,0.1, p = 0.0014; FC-TTX:0.13,0.01,0.21, p = 0.16). The median occurrence of isolated spikes in the No-TTX and the CH-TTX groups, however lower in the latter, was not significantly different (No-TTX: 3.84,2.41,5.89; EF-TTX:0, p<0.0001; CH-TTX:1.11,0.35,4.56, p =0.11; FC-TTX:2.43,1.86,3.94, p=0.48).

### Silencing the contralateral hippocampus suppresses HPDs but not interictal epileptic activities

In the same group of mice, we then silenced selected regions using TTX, at d30: the focus region (n=10 mice), the contralateral hippocampus (n=11) or the ipsilateral frontal cortex (n=10). As expected, silencing the focus completely suppressed focus epileptiform discharges (Fig.2C). Contralateral hippocampus silencing also completely abolished both subtypes of HPDs (sHPDs and iHPDs) in all mice, as efficiently as focus silencing. Conversely, STs and ISs were still generated in the focus during contralateral hippocampus silencing, though the number of STs was reduced. TTX injections in the ipsilateral frontal cortex did not impact the expression of any epileptiform discharges, supporting a specificity of the contralateral hippocampus in affecting focus epileptiform discharges.

It is crucial to note that, as previously described ^4^, injecting a small volume of TTX decreases background EEG activities above the injected region but not in other regions. Accordingly, iEEG recordings show that TTX injections in the contralateral hippocampus reduces neuronal activities and background power in the injected region but not in the focus (Fig.S3), confirming the localized effects of TTX.

### Impact of region-specific silencing on the expression of epileptiform discharges in the epileptic network

Beside focus epileptic activities, interictal large-scale network IEDs also characterizes the kainate model, reflecting the development of a large-scale network of epileptic regions. Using HD-EEG, and confirming our previous results ^4^, silencing the focus did not reduce the rate of network IEDs (Fig.3A, C). Silencing the ipsilateral frontal cortex region did not decrease network IEDs either, but silencing the contralateral hippocampus was associated with a significant reduction.

**Figure 3.**
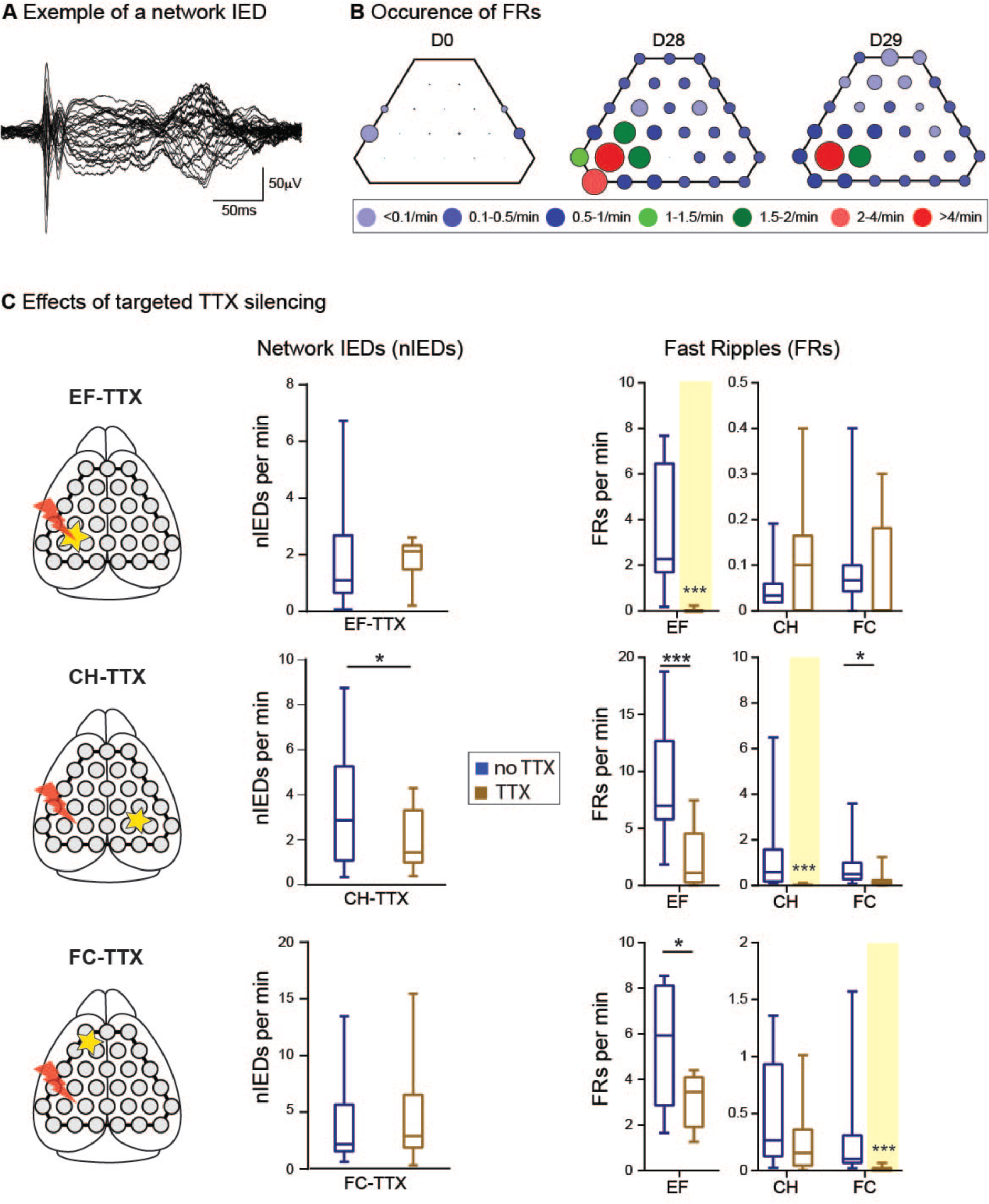
Silencing the contralateral hippocampus reduces the occurrence of epileptiform discharges in the epileptic network. A) Illustrative example of network IEDs (averaged raw LFP of network IEDs from one recording session) recorded with HD-EEG. The superimposed average traces illustrate the typical spike and waves pattern invading all surface electrodes. B) Localization and occurrence of FRs in mice recorded at D0 and during the chronic stage (D28, D29; same groups as in Fig.2B) illustrating the absence of FRs at D0, with the exception of the somatosensory cortex as previously demonstrated (see supplementary information), and the increased expression of FRs at the large-scale level (D0: 0.4, 0.15, 0.78; D28: 12.63, 5.83, 24.5, p=0.0001; D29: 12.92, 4.17, 19.11, p=0.01(vs D0), p=0.39 (vs D29), unpaired one-way ANOVA). Within each color-coded classes of occurrence rate, the size of the dot varies in function of the rate values. C) Effects of targeted TTX silencing on the expression of interictal network IEDs and FRs. The left box plots show the occurrence of network IEDs in chronic epileptic mice recorded before and during TTX silencing in the epileptic focus (EF; median and IQ range: No TTX: 1.16,0.34,2.63; TTX: 2.1,1.09,2.25, p =0.42, n = 10), the contralateral hippocampus (CH; No TTX: 2.94, 1.1, 5.42; TTX: 1.28, 0.8, 3.26, p = 0.01, n = 11) and the ipsilateral frontal cortex (FC; No TTX: 2.33, 1.68, 5.94; TTX: 2.90, 1.84, 6.67, p = 0.37, n = 10, Wilcoxon tests). Note that the network IED and wave pattern was grossly similar, even during targeted TTX silencing although the electrodes above the silenced regions did not exhibit the epileptic discharges, as expected (see supplementary figure S4 for illustrations of network IEDs waveforms in all groups). The right box plots show the occurrence of interictal FRs (excluding FRs recorded during HPDs and network IEDs) before (no-TTX, blue) and during (TTX, brown) TTX silencing detected at electrodes above selected regions of interest (EF, CH, FC). In all cases, FRs were completely abolished in the silenced region, highlighted in yellow. The effects of TTX silencing of one region upon FR expression in other ROIs are described in the main text (EF silencing: in CH, No-TTX= 0.03, 0.01, 0.06, TTX= 0.1, 0, 0.16, p = 0.25 and in FC, No-TTX: 0.06, 0.04, 0.1; TTX: 0, 0, 0.18, p = 0.25; CH Silencing: in EF, No-TTX= 6.98, 5.73, 12.74; TTX= 1.12, 0.2, 4.63; p = 0.001 and in FC, No-TTX: 0.49, 0.23, 1.02; TTX: 0.13, 0.03, 0.26; p = 0.03; FC silencing: in EF, No-TTX: 5.92, 2.83, 8.14; TTX: 3.44, 1.88, 4.12, p = 0.01 and in CH, No-TTX: 0.26, 0.11, 0.93; TTX: 0.15, 0.03, 0.36, p = 0.62). EF= epileptic focus; CH= contralateral hippocampus, FC= frontal cortex.

We next measured the impact of targeted silencing on the expression of interictal FRs, another epileptic biomarker ^25,26^. Interictal FRs were defined as those non-concomitants with HPDs and network IEDs, as both events were shown to be reduced with contralateral hippocampus silencing. As previously shown ^4^, we detected interictal FRs across both hemispheres during the chronic stage of the disease (D28-29), with a particularly high incidence rate in the focus region (Fig.3B). Targeted silencing affected differently interictal FRs depending on the silenced region. Focus silencing did not reduce the occurrence of remote FRs (Fig.3C). Silencing the frontal cortex induced a small reduction of interictal FRs in the focus but not in the contralateral hippocampus. Conversely, contralateral hippocampus silencing led to a strong and significant reduction of interictal FRs in the 2 other regions of interest, i.e. focus (i.e., ipsilateral hippocampus) and frontal cortex, further highlighting the major role of the contralateral hippocampus node in the large-scale epileptic network.

### Intracortical EEG confirms that contralateral hippocampus silencing abolishes paroxysmal discharges but not interictal epileptic activities

Our results show that silencing the contralateral hippocampus prevents the focus from generating paroxysmal discharges, including iHPDs reminiscent of focal seizures, but not interictal epileptic spikes (IS and ST). HPDs are recorded throughout the DG and the cornus amonis with phase inversions indicating the location of generators in the DG ^4^ (Fig.S2). To investigate further the role of the contralateral hippocampus in the generation of focus epileptiform activities, we used bi-hippocampal linear probes spanning the dorsal hippocampus at the level of the focus and the homologous region of the contralateral hippocampus.

Figure 1 shows illustrative examples of the 4 classes of epileptiform discharges recorded with iEEG in the epileptic focus in 16 mice. Confirming HD-EEG quantifications, isolated spikes constituted the majority of detected events with a median occurrence rate of 3.6/minute (Fig4.B). Spike trains had a median occurrence rate of 0.68/minute and a mean±SD duration of 2.05±1.72 sec. Ictal HPDs were observed in 11 of 16 mice, at a rate of 0.13/minute, and a duration of 27.12±16.16 sec. Short HPDs were less frequent and had a duration of 5.56±2.64 sec. As illustrated in Figure 1D, all HPDs started with a low-frequency train of several typical high amplitude spikes and then gradually escalate to high frequency spiking, which fits with the hypersynchronous onset seizure type predominantly reported in the kainate model ^13,23,27^.

Silencing the contralateral hippocampus fully abolished all HPDs in all tested animals (n=8 TTX; Fig.4B). STs were reduced but not abolished and their duration remained stable, at about 2 sec. The occurrence of ISs remained unaltered, confirming HD-EEG results. Furthermore, ISs retained similar intrinsic parameters characteristics than before silencing in terms of spike amplitude, peak oscillatory frequency, duration, and number of oscillations of the associated FRs (Fig.4C). This indicates that contralateral hippocampus inhibition specifically impacts on paroxysmal activities of the focus without altering the morphology of isolated epileptic discharges. Altogether, our results suggest a role of contralateral hippocampus activity in the transition from states marked by short interictal to longer epileptic activities. In the following section, we thus wanted to investigate the potential mechanisms underlying the association between contralateral hippocampus activity and the expression of longer-lasting epileptic activities in the focus.

**Figure 4.**
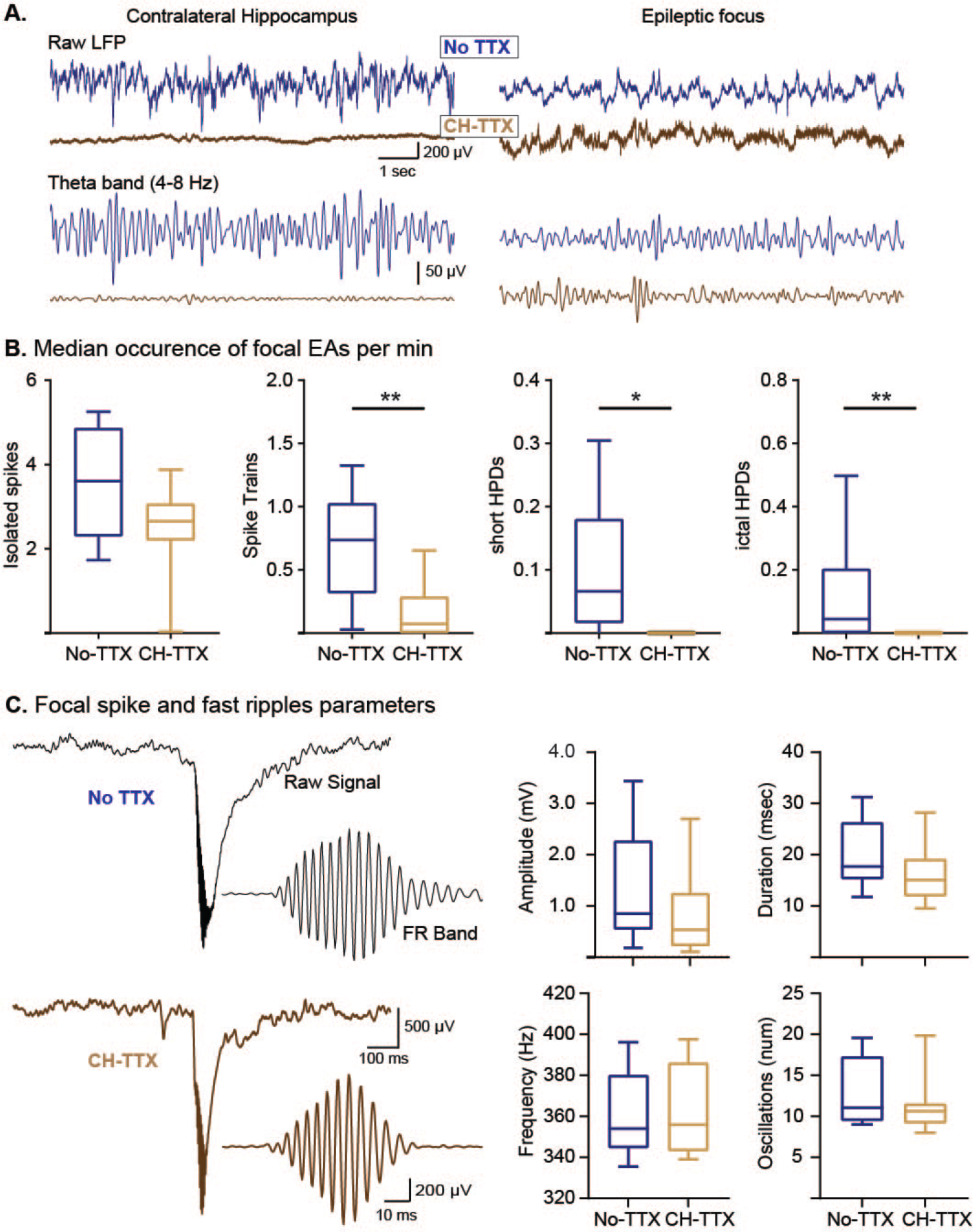
Contralateral hippocampus (CH) silencing reduced spike trains and abolished HPDs during iEEG recordings. A) Illustrative traces of background raw LFP and corresponding theta filtered signal recorded simultaneously in the focus and the CH before and during TTX injection in the CH (no TTX vs TTX) using iEEG. TTX efficiently and specifically suppresses LFP activities locally in the CH but not in the focus (see supplementary Fig.4 for background LFP quantifications). B) Median (with IQ range and 10-90 percentiles) occurrence of focal EAs (No TTX=16, CH TTX=8): isolated spikes: No-TTX: 3.60, 2.3, 4.86; CH-TTX: 2.65, 2.19, 3.06, p = 0.09; Spike trains: No-TTX: 0.73,0.31,1.02; CH-TTX: 0.07,0,0.28; p = 0.004; short HPD: No-TTX: 0.066,0.01,0.18; CH-TTX: 0; p = 0.04) and ictal HPD: No-TTX: 0.04, 0, 0.2; CH-TTX: 0, p = 0.0035 (Mann Whitney non-parametric tests). C) Illustrative traces of isolated spikes and overriding FRs in the focus during no TTX and CH TTX recordings (raw LFP). The filtered signals in the FR band (200-550 Hz) are shown next to the respective ISs. CH Silencing did not alter the amplitude of the focal isolated spikes (peak amplitude in the 1-40 Hz band) nor the morphological characteristics (frequency, duration, number of oscillations) of the overriding fast ripple. Whisker plots histograms show the quantifications of FR parameters in no TTX and TTX recordings (Median, IQ, 10-90 percentiles).

### Contralateral hippocampus MUA anticipates focus epileptiform discharges

Neuronal to slow oscillations coupling has been suggested to be involved in the generation of epileptic activities ^28,29^. We tested whether such coupling could be involved in the presumed role of the contralateral hippocampus in dictating the transition between interictal to ictal states.

Figure 5A shows a typical IS, with the characteristic phase inversion centred on the molecular layer of the DG. The raw signal is overlapped with the MUA detected from the corresponding contralateral electrodes, revealing MUA bursts starting in the contralateral hippocampus around 200 ms before the spike and the associated MUA increase in the focus. We then systematically quantified MUA in the focus and the contralateral hippocampus in the periods preceding focus epileptiform discharges and compared it to MUA rates recorded during backgrounds periods. We analysed separately periods preceding ISs from periods preceding STs, sHPDs and iHPDs combined (=ST+HPDs) since contralateral hippocampus silencing specifically impacted these more intense epileptiform discharge types, i.e., events of longer duration and higher spike loads. This analysis showed that contralateral MUA rate was significantly higher prior to epileptiform discharges as compared to background (Fig.5B). Furthermore, contralateral MUA bursts started earlier than MUA in the focus (Fig.5C) which, incidentally, increased significantly only before ST+HPDs but not before IS (Fig.5B). Importantly, the increases in contralateral MUA also significantly preceded the LFP increase associated with the epileptiform discharges in the focus, i.e., the presumed onset of IEDs. Altogether, contralateral MUA starts before any other recognizable activity associated with ipsilateral EA.

**Figure 5.**
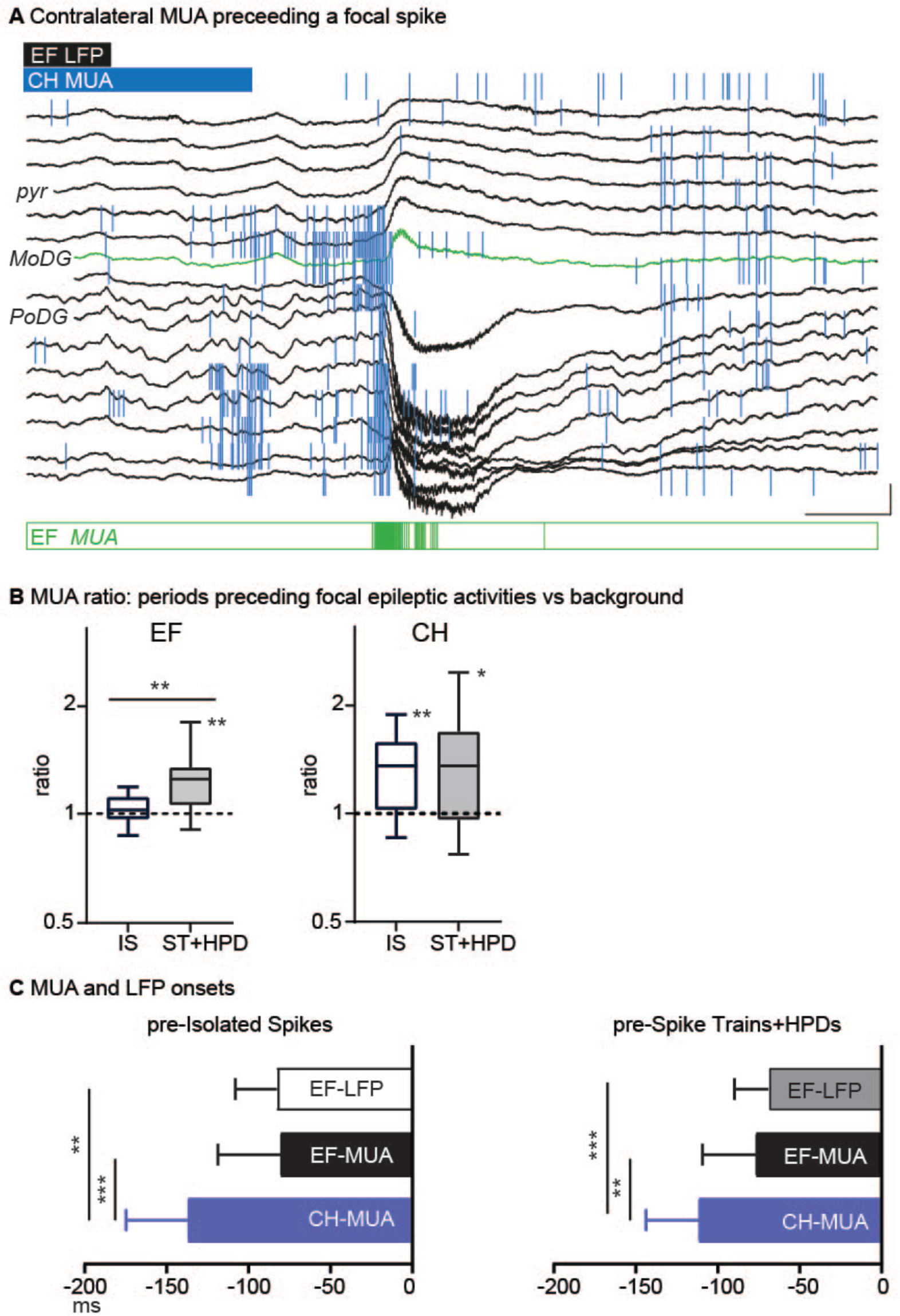
MUA in the CH precedes focal epileptic activities. A) Example of a focal epileptic discharge (black, raw LFP) overlapped with the contralateral action potentials (MUA activity, blue vertical lines), illustrating the strong MUA activity in the contralateral hippocampus (CH) before the epileptic discharge in the EF. Below, MUA activity in the epileptic focus (EF) as measured from the channel highlighted in green. Scale: 2 mV, 100 ms. B) Median (+IQ range and 10-90 percentiles) ratios of MUA per second in the 1 sec-periods preceding the statistical onset of epileptic activities as compared to the background (periods devoid of any epileptic activity) in the EF (IS: 1.04, 0.88, 1.47; ST+HPDs: 1.39, 1.14, 2.04) and the CH (IS: 1.18, 0.99, 1.44; ST+HPDs: 1.41, 0.93, 1.71) across mice. In the EF, MUA was significantly higher in the pre-EA periods than during background periods for ST+HPD (median increase: 24.5 %, p <0.01, Wilcoxon test). The MUA ratio was also significantly higher before ST+HPD as compared to pre-ISs periods (p=0.01). In the CH, the MUA expression was significantly stronger before both ISs (median increase: 35.5%, p<0.01) and ST+HPDs (median increase: 35.6%, p<0.05) as compared to the background periods. C) Median (+IQ range) of statistical onsets of LFP in the EF (IS and ST+HPD) and of MUA in the EF and CH (relative to the GFP peak of the epileptic discharge, see methods) across mice. MUA in the CH started significantly before both MUA (median difference of 54 ms for IS and 42 ms for ST+HPD) and LFP (median difference of 48 ms for IS and 43 ms for ST+HPD **p<0.01, ***p<0.001) in the EF.

### Delta/theta band power increases in both hippocampi before focus epileptiform discharges and more strongly ahead of more intense epileptic discharges

What could be the nature of the interaction between the focus and the contralateral hippocampus region, seemingly involved in the generation of focus epileptic activities and in particular long-lasting ones? To address this, we analysed the functional interactions between the focus and the contralateral hippocampus in the periods preceding epileptiform discharges. The illustrative spectrograms in figure 6A show peaks in the delta (.5-4 Hz) and theta (4-12 Hz) range in periods preceding focus epileptic activities, although the theta peak is less marked in the EF as previously observed in the model ^30^. Quantitative analyses showed that delta and theta band amplitudes increased significantly before all epileptiform discharges in both regions as compared to background (Fig.6B), except before IS in the contralateral hippocampus. The increase in amplitude in the theta band is stronger for periods preceding ST+HPDs than before IS in the focus (p=0.0012) and the contralateral hippocampus (p=0.006).

**Figure 6.**
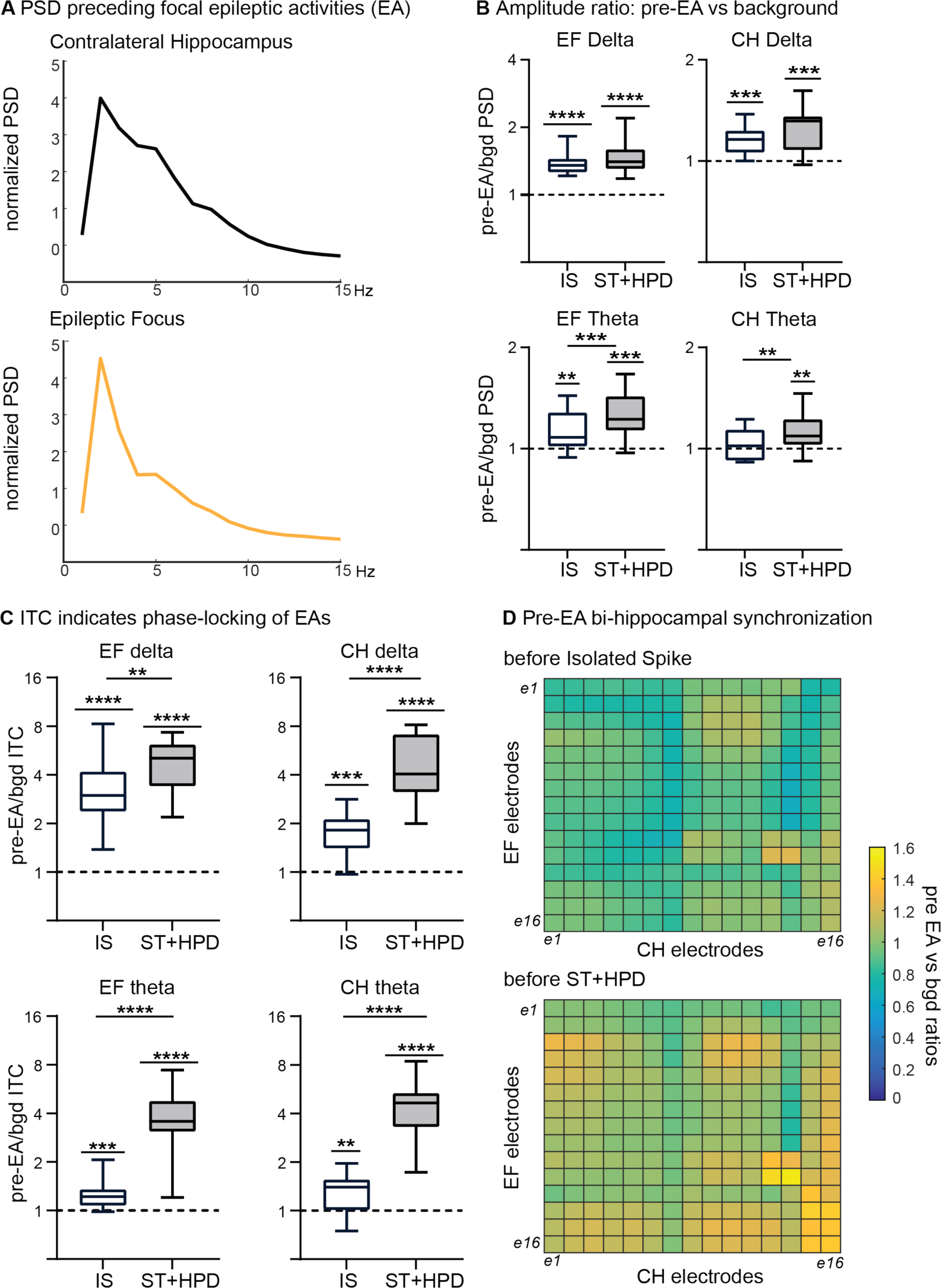
Increased delta/theta amplitude and bi-hippocampal synchronization before focal epileptic activities (EA) during iEEG recordings. A) Illustrative example of normalized PSD (z-scored normalized against background periods free of IED) from one mouse and all EA types over the pre-EA period (−1700 to −700 ms before EA first peak). B) Amplitude ratios in delta and theta frequency bands before EAs as compared to background (bgd). Amplitude increased in EF and CH in both delta (median increase: EF: IS=35%, p <0.001; ST+HPDs=40%, p<0.001; CH: IS=16%, p =0.0006; ED=31%, p =0.0009, one-sample Wilcoxon test) and theta bands (median increase: EF: IS=8%, p =0.01; ST+HPDs=22%, p=0.0027; CH: IS=2%, p =0.62; ED=9%, p =0.02). C) ITC during pre-EAs periods increased significantly as compared to background in the delta and theta frequency bands both in EF (median increase: IS=197%, p <0.001 and ST+HPDs=414%, p<0.001 for delta band; IS=21%, p <0.001 and ST+HPDs=273%, p<0.001 for theta band; one sample Wilcoxon test with Bonferroni correction) and CH (median increase: IS=81%, p =0.003 and ST+HPDs=304%, p<0.001 for delta band; IS=39%, p =0.012 and ST+HPDs=384%, p<0.001 for theta band) indicating that epileptic activities are phased locked with slow oscillations in delta and theta frequency bands in both regions. ITC was stronger before ST+HPDs than before IS (p values for comparisons between IS and ST+HPDs: focus=0.0036 and <0.001 for delta and theta; contralateral hippocampus <0.001 for both delta and theta). D) Theta band synchronization maps between CH and EF from one animal, illustrating the increase in synchronization between EF and CH before focal epileptic activities. Note the stronger increase in bi-hippocampal synchronization before the occurrence of HPDs and spike trains than before isolated spikes (warmer colors in the below graph).

### Phase-locking of focus discharges to the delta/theta bands of the focus and the contralateral hippocampus

Given the increase in delta/theta power in the bi-hippocampal network prior to epileptic events in the focus, we investigated phase-locking of these epileptiform discharges to delta/theta bands in the focus and the contralateral hippocampus. We estimated phase-locking by comparing inter-trial coherence (ITC) between pre-epileptiform discharges and background periods. ITC was significantly higher before IS and ST+HPDs than in background epochs for both delta and theta frequency ranges, and these increases were significantly stronger before longer-lasting epileptiform discharges, i.e., ST+HPDs, than before IS (Fig.6C). These analyses indicate that focus epileptiform discharges are phase-locked to delta/theta bands of both hippocampi, suggesting a synchronization of the bi-hippocampal network prior to those events.

### Bi-hippocampal theta synchronization increases before spike trains and HPDs

Accordingly, we tested whether phase-synchronization between the focus and the contralateral hippocampus increases before epileptiform discharges by calculating the ITC of the phase differences between contacts of both hippocampi. Figure 6D shows, for one illustrative animal, the theta band synchronization maps of all electrode pairs between the 2 hippocampal probes (pre-epileptiform discharges ITC normalized by baseline), illustrating a general increase in the ITC ratios before ST+HPD. Across all animals, phase-synchronization in theta increased before ST+HPDs events by 15% (median and IQ range: 1.15, 1.01, 1.31; p=0.0078) while no such effect was noticed before IS (1.009, 0.93, 1.06, p>0.99). In agreement with that, phase-synchronization was significantly stronger for ST+HPDs than for IS (p=0.0012).

## Discussion

Here, we revealed that the bi-hippocampal network is crucial for ictogenesis. Contralateral hippocampus silencing is associated with a suppression of long-lasting epileptic activities (HPDs), a decrease of intermediate epileptic discharges (ST) and no effect on isolated spikes. Altogether, this suggests a predominant role of the contralateral hippocampus in the transition from interictal to ictal – or at least longer-lasting – activities. Transitions from highly focal “microseizures” to “macroseizures” have been identified in humans ^10^. Although the comparison between these microseizures and the pathological events that we describe is challenging, the identification of the mechanisms that allow such transition is crucially important. In addition, we observed that the contralateral hippocampus is part of a large-scale irritative network in which IEDs are produced, i.e., network IEDs and remote FRs, and plays a dominant role in this network since its pharmacological silencing, unlike focus silencing, reduces the production of IEDs in the other nodes of the network.

### A bi-hippocampal epileptogenic network

What mechanisms explain the selective impact of contralateral hippocampus silencing on focus paroxysmal activities but not isolated spikes? Neuronal activity, as measured with MUA and PSD analyses, increases both in the contralateral hippocampus and the focus prior to epileptiform discharges, and more strongly ahead of longer and more intense activities, i.e., STs and HPDs. A striking observation comes from the strong increase in ITC in the delta/theta range prior to STs and HPDs. The fact that the increase in ITC is observed for signals from both hippocampi reflects the bi-hippocampal synchronization within this low frequency range that specifically rises in the period preceding STs and HPDs. The 2 hippocampi are reciprocally connected ^31,32^ and silencing the contralateral hippocampus, by reducing its inputs towards the focus, might decrease the level of activity in the latter and therefore the chance of overpassing the seizure threshold. More specifically, one could hypothesize that the build-up of focus excitation required for the transition from interictal to ictal states is aborted as a consequence of reduced synchronization in the bi-hippocampal network following contralateral hippocampus silencing, and consequently the number of STs and HPDs as well. Theta activity can still be observed in the epileptic kainate-injected hippocampus, although it is significantly decreased, in comparison to the contralateral hippocampus ^30^. Previous studies have shown an increase in theta power in the pre-ictal state ^33^ or in the period immediately preceding IEDs ^28^. Still, the involvement of theta rhythms in epilepsy has proven to be complex, intervening in different ways depending on the etiology of the disease, the seizure phase as well as on the location in the seizure-onset-or propagation-zone ^34–37^.

These results are in line with current knowledge on the contribution of large-scale interactions in the development of abnormal brain activity in epilepsy. For example, interactions between the focus and basal ganglia are involved in seizure modulation in patients and animal models ^38,39^. Furthermore, in the kainate model, optogenetic modulation of distant neurons, including in the contralateral hippocampus, can decrease duration or propagation of ongoing seizures ^2,32,40,41^. Another example comes from network modelization studies, which suggest that large-scale connectivity changes in epilepsy better explain the emergence of seizures than purely local mechanisms ^11,42^.

### The transition from spikes to paroxysmal discharges

Typically, HPDs start with a period of high-amplitude low frequency spikes before escalating to higher spiking rates, which correspond to the hypersynchronous seizure onset type in kainate models ^13,23,27^. Spikes at HPDs onset are closely similar to those characterizing IS and ST, suggesting that HPDs could be initiated by the same neural networks as those producing transient interictal spikes, as already previously hypothesized ^13^. When spikes occur during a period of strong synchronized activity in the bi-hippocampal network, they could trigger a phase of recurrent excitation likely to evolve into an ictal phase. The specific suppression of HPDs upon silencing of the contralateral hippocampus supports this hypothesis, in line with recent findings reporting that artificially synchronizing the activity of the two hippocampi promotes seizure generation ^43^. Our findings only posit that contralateral hippocampus silencing is associated with an interictal state where long-lasting epileptic events are prevented, rather than establishing that the contralateral hippocampus prevents isolated spikes to progress into longer-lasting epileptic activities, such as HPDs. That HPDs appear as spikes escalating towards ictal discharges in particular conditions does not mean that IEDs are aborted seizures. IEDs are clinically relevant as markers of the seizure-onset zone ^44^ and have been involved in transient cognitive impairments including memory disturbances ^45,46^. The link between IEDs and seizures is highly debated; increases in IEDs in the preictal period have been proposed to either favor the generation of seizures or prevent them to occur ^43,47,48^. Still, IEDs could reflect interictal actvity of seizure-generatng networks ^47,49^ and therefore investgatng the modulaton of these events is important to understand seizure initaton and develop novel therapies aimed at decreasing the risk of transiton towards more debilitatng epileptc events ^2,3^.

## Conclusions

This study provides evidence that state transition from shorter to more sustained epileptic activities is, at least in part, governed by a large-scale network where the contralateral hippocampus plays a decisive role. Cross-talk between contralateral neuronal activity and ipsilateral delta/theta oscillations could be a mechanism by which the contralateral hippocampus favors sustained epileptic activities. While extra-focal pathological activities are suspected to be major components of the disease, we know little about the large-scale network mechanisms that govern transitions to ictal activities. Altogether, this adds to the growing body of evidence that monitoring the activities from several nodes and acquiring a thorough understanding of their interactions is crucial to develop more sensitive closed-loop devices aimed to control seizures.

## Acknowledgments

GPP was supported by the Swiss National Science Foundation (grant n° 140332). CM was supported by the Swiss National Science Foundation (grant n° 320030-159705), by the National Center of Competence in Research (NCCR) “SYNAPSY”, and by the Center for Biomedical Imaging (CIBM) from Geneva and Lausanne. LS was supported by the Swiss National Science Foundation (grant n° 323530-158125 and P500PM_206720). CQ was supported by the Swiss League Against Epilepsy.

## Author contributions

GPP and FF: Acquisition and analysis of data, drafting a significant portion of the manuscript and figures. KS: Conception and design of the study. CMM : Conception and design of the study, analysis of the data. LS: Conception and design of the study, analysis of data, drafting a significant portion of the manuscript. CQ: Conception and design of the study, analysis of data, writing manuscript and figures.

---

**Supplementary Figure S1.**
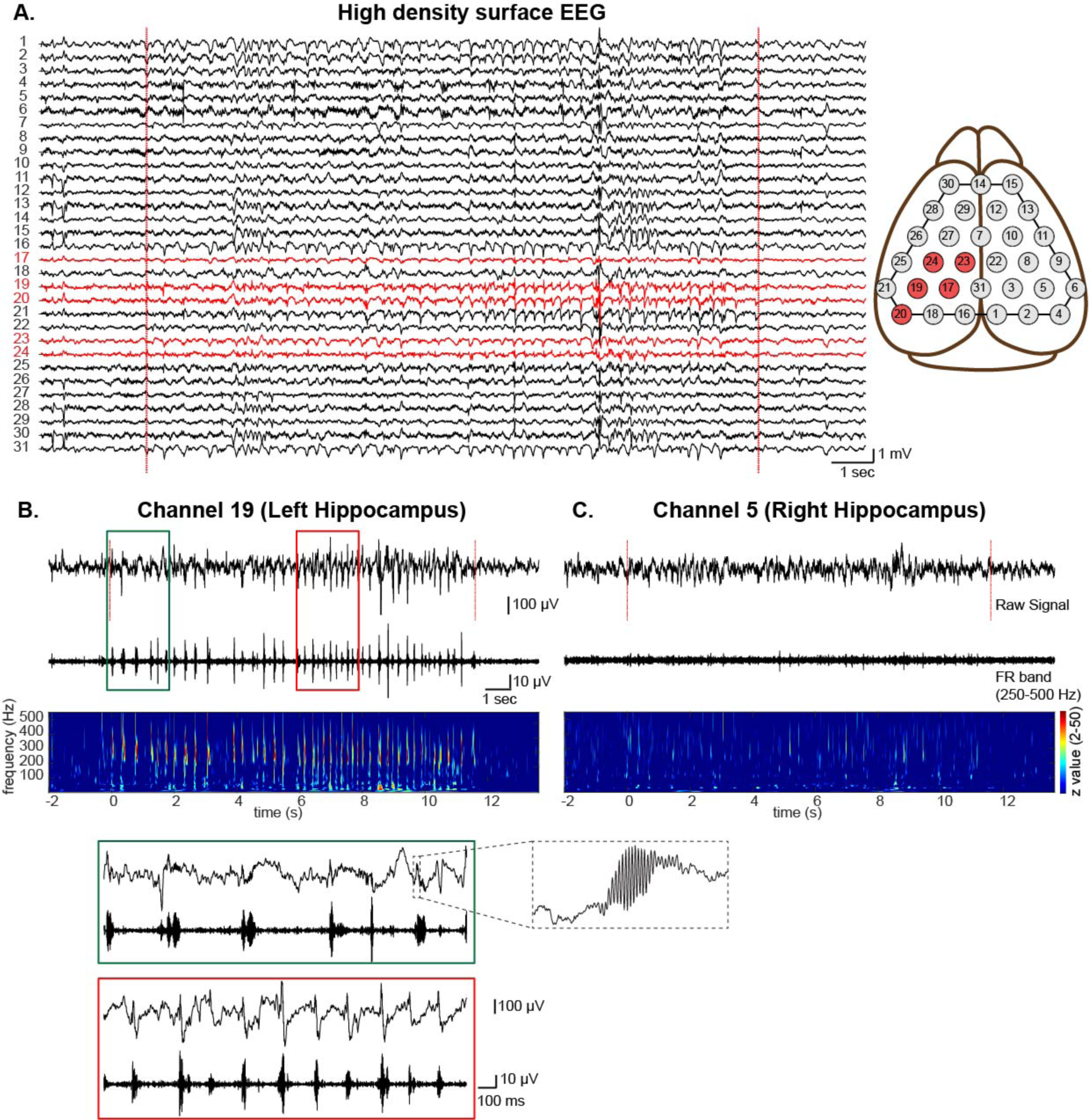
Illustration of awake head-fix epileptic recording with HD-EEG. A) Raw signal illustrating a typical hippocampal paroxysmal discharge (HPD) with rhythmic spikes reminiscent of a focal seizure, lasting 16 sec (red dotted lines indicate starting and end points of the event) detected at electrodes above the kainate injected hippocampus (red traces). The approximate positions of the surface electrodes are illustrated in the right cartoon. B, C) Zoom on the signals recorded at single channels from the left (kainate-injected) and right (contralateral) hippocampus, respectively. Top panels: raw LFP; middle panels: filtered signal in the FR band (250-550Hz). Bottom panels: normalized time-frequency plot across the 0-500 Hz spectrum showing increased activity in the spike (10-40 Hz) and FRs band at the electrode above the kainate injected hippocampus. The green and red insets represent expanded 2 sec views of the onset and middle periods of the HPD, illustrating a progressive increased in spiking rates. *FRs are readily visible in the raw LFP signal, as illustrated by the zoom in the dotted inset*.

**Supplementary Figure S2.**
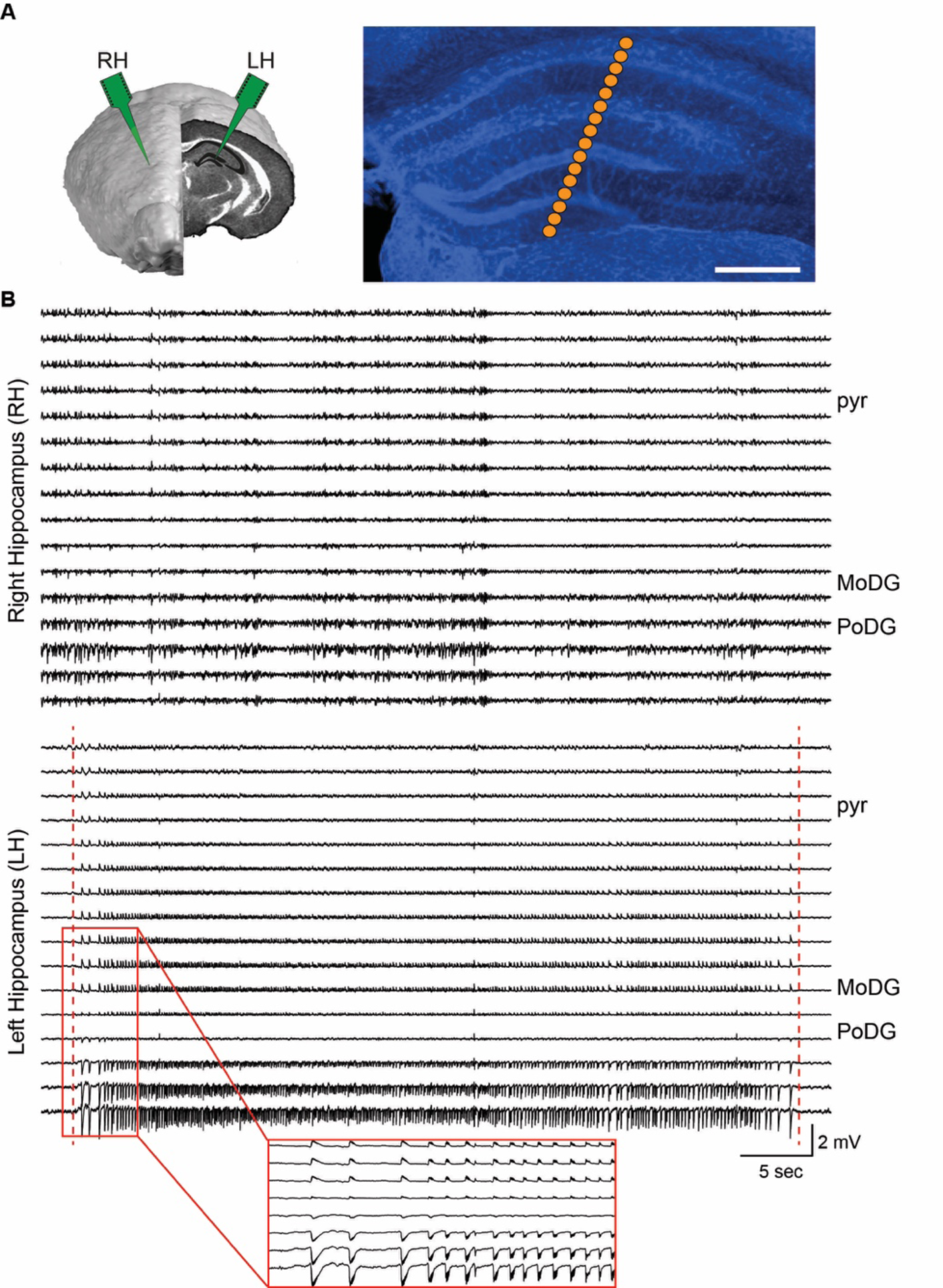
Intracerebral bi-hippocampal head-fixed recording (iEEG) in chronic epileptic mice during a focal seizure. A) Two A16 (16 channels) neuronexus probes were inserted into the left (kainate-injected, LH) and right hippocampus (RH). The DAPI stained picture illustrates the targeted positions of the 16 channels. Scale bar=500 (m. B) Intracerebral recording of a ictal HPD lasting 46 seconds, reminiscent of a focal seizure. The red lines denote the onset and offset of the seizure. Discharges show clear phase inversions at the level of the dentate gyrus in the injected hippocampus (LH). Inset: 5 seconds zoom of the focal seizure start suggesting a hypersynchronous onset seizure type. Approximate positions of the molecular layer of the dentate gyrus (MoDG, outer shell), the polymorph layer of the dentate gyrus (PoDG, hilus) and of the pyramidal layer of CA1(pyr) are indicated.

**Supplementary Figure S3.**
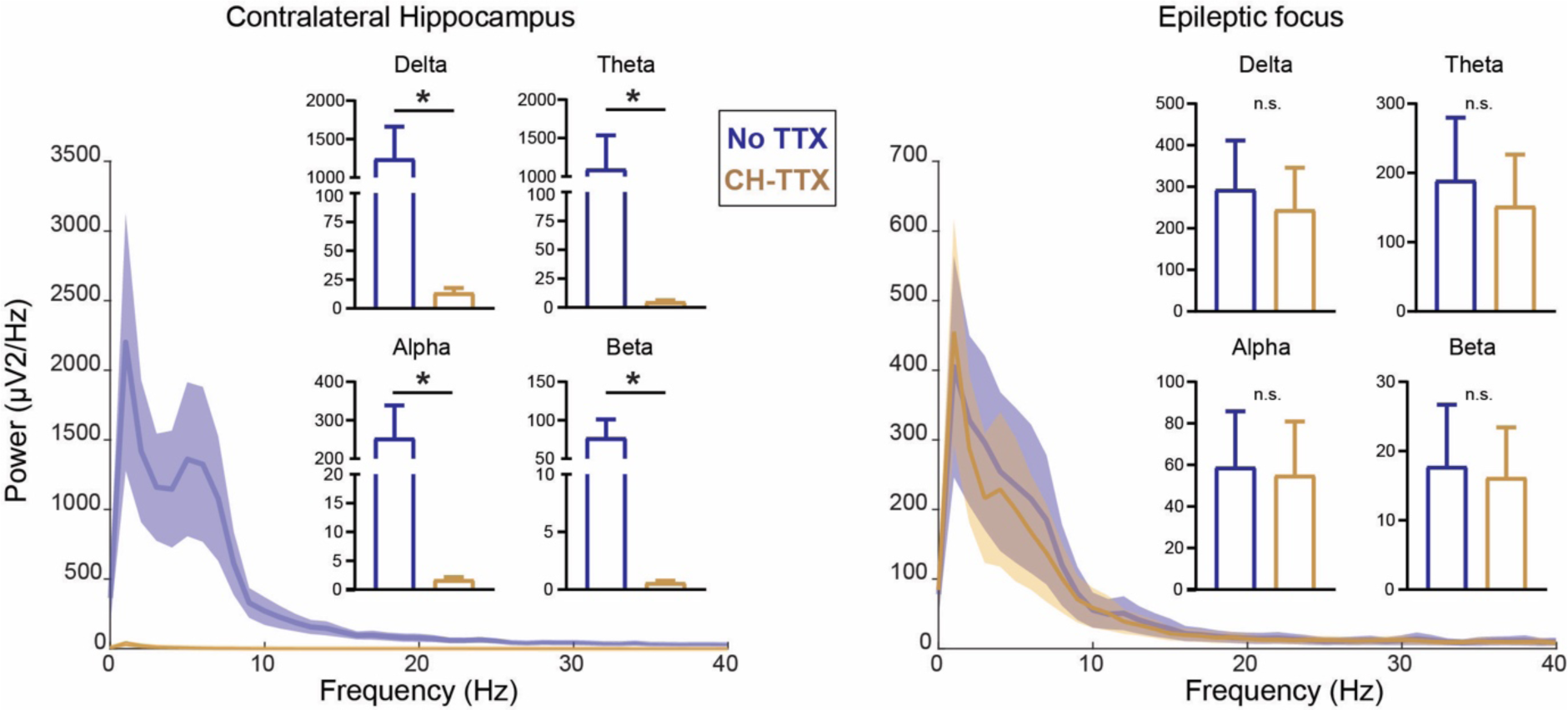
TTX silencing of the CH does not reduce background activity power in the EF. A 2-way ANOVA with factor 1 as silencing and factor 2 as frequency range revealed that there was an effect of silencing in the contralateral hippocampus (silenced region, CH) but not in the epileptic focus. We then performed a non-parametric individual t-test between No-TTX and CH-TTX in in different frequency bands and corrected for multiple comparisons. Mean±SEM background power spectral density in CH and EF before and during silencing across all mice (n=8). Background power significantly decreased during the TTX injection in the contralateral hippocampus in all frequency bands as compared to the No TTX sessions at p<0.05 (Mean±SEM; delta: No-TTX: 1242.8±422.2, TTX: 13.44±4.35; theta: No-TTX: 1094.8±439.9, TTX: 4.66±1.464; alpha: No-TTX: 253.4±85.48, TTX: 1.69±0.48, beta: No-TTX: 77.06±24.16, TTX: 0.5±0.17). Background power remained unaltered in the EF during silencing of the CH. Epileptic focus (Mean±SEM; delta: No-TTX: 292.6±118.6, TTX: 243.9±101.7; theta: No-TTX: 188.8±90.94 TTX: 151.5±74.75; alpha: No-TTX: 58.89±26.98, TTX: 54.97±25.97; beta: No-TTX: 17.77±8.94, TTX: 16.17±7.27, all p >0.05). Delta: 0.5-4 Hz; Theta: 4-8 Hz; Alpha: 8-12 Hz, Beta: 12-30 Hz.

**Supplementary figure S4.**
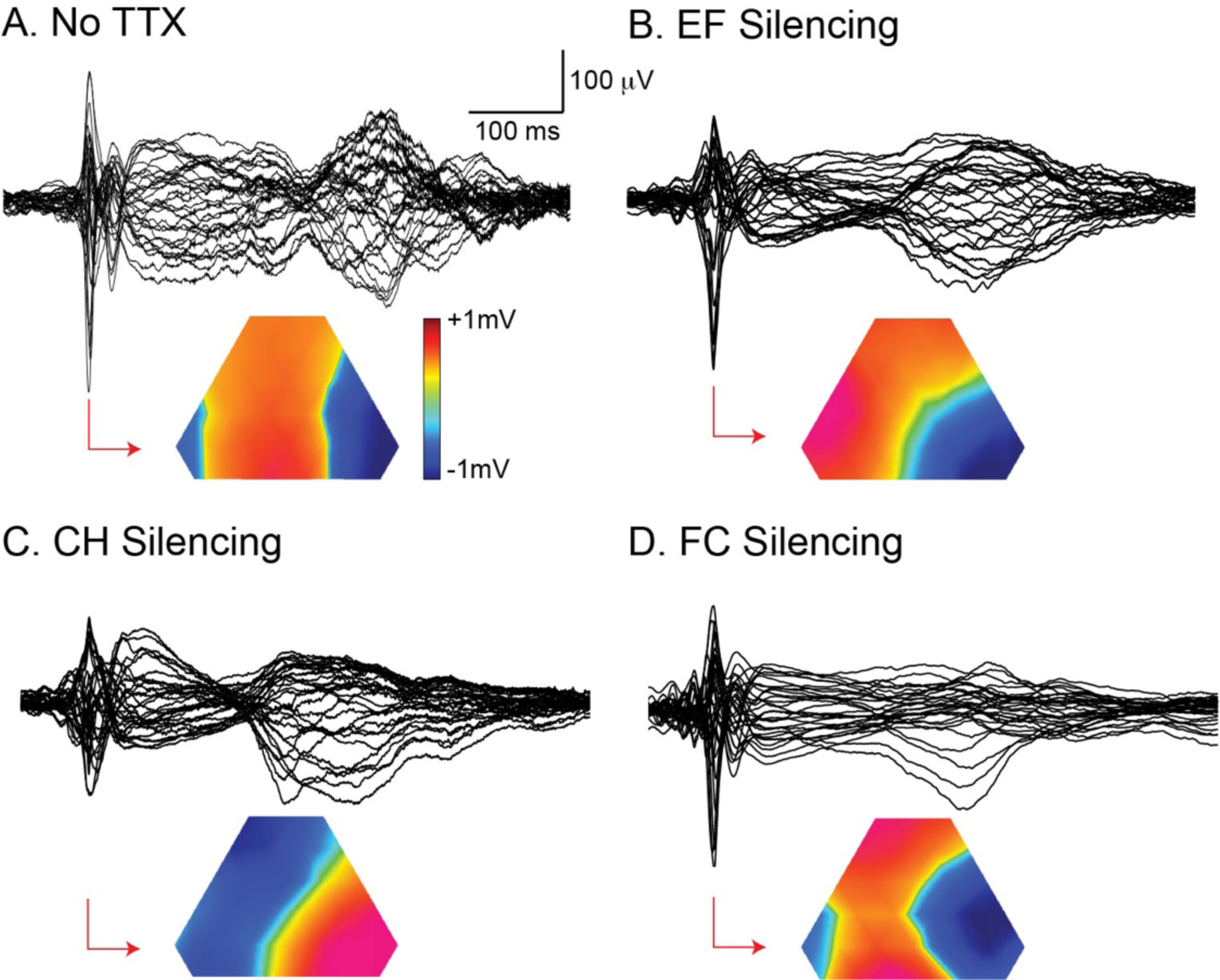
Examples of network IEDs recorded with HD-EEG in epileptic animals. The superimposed traces (average raw LFPs at each electrode from one recording session) illustrate the typical spike and wave patterns invading all surface electrode during recording made without TTX injection and during TTX silencing in the epileptic focus (EF), the contralateral hippocampus (CH) and the ipsilateral frontal cortex (FC). Below the network IEDs traces, the interpolated voltage maps obtained at the initial peak of the IEDs are shown. The topography of the onset maps in the NO TTX and FC silencing recordings displays the lateral negative and medial positive pole thant flanked the positions of the 2 hippocampal axes and reflect the activity of concomitant sources across both hippocampi, as previoiusly demonstrated (Sheybani et al, 2018). Note that in our earlier study, network IEDs were coined “generalized spikes”. As expected, the TTX injected hippocampus dipole is absent in the EF and CH silencing recordings.

## Supporting Materials and Methods

### Animals

Forty-seven male C57BL/6j adult mice (Charles River), all around 10–11 weeks old by the time they entered the protocol, were included in the study. Thirty-one of these mice were recorded using surface high-density EEG (HD-EEG) at day 28 (d28) post-kainate injection, of which 15 were recorded at d-1 before kainate injection as controls and 20 also on a second day during the chronic stage, at d29 post-kainate. TTX silencing experiments (see below) were performed on d30 or d31, with TTX injections in either the EF (n=10), the contralateral hippocampus (n=11), an important source of bilateral homologous connections (Bui et al., 2018) or the ipsilateral frontal cortex region (primary motor cortex; n=10), a region that produce FRs during the chronic stage (Sheybani et al., 2018). The intracortical data comes from a total of 16 mice that were recorded with bi-hippocampal intracortical EEG (iEEG) on d28 post kainate injection. Eight of these mice came from an earlier study (Sheybani et al., 2018, 2019). In the 8 other mice, recordings were made on d28 and then further during contralateral hippocampus silencing on d29 or d30. All animal procedures were performed in accordance with the Geneva and Switzerland animal care committee’s regulations.

### Surgeries for surface awake head-fixed HD-EEG and iEEG recordings

Head-fix surgeries were performed under injectable anesthesia (Medetomidine (Domitor): 0.5 mg/kg, Midazolam (Dormicum): 5 mg/kg, Fentanyl: 0.05 mg/kg) injected intra-peritoneally (i.p.). Animals were mounted on a stereotaxic frame and an aluminium ring-like header attached to the mouse skull using dental cement as previously described (Sheybani et al., 2018). The skin was retracted using four bulldog clamps to keep the incision open, and the conjunctive tissue removed using a surgical blade and cleaned with sterile saline solution. For surface EEG recordings, the positions of the EEG electrodes were marked by staining the grid of 32 stainless steel electrodes using blue ink and placing them over the skull. A drop of Loctite (Henkel) was spread over the skull to form a thin layer and once the Loctite is dry, a small patch (500µm) of the Loctite was removed by drilling over the electrode marked positions using a nano drill (World Precision Instruments). For bi-hippocampal recordings, the positions of the future craniotomies for neuronexus probes acute implantations were marked with blue ink before covering the skull with Loctite glue. For surface and intracortical recordings, the site of injections for kainate and tetrodotoxin was also marked with ink. The centre of the aluminium ring-like holder was then filled with protective silicon (Kwik-Cast, World Precision Instrument), which can be easily removed and replaced to allow easy access for further injections and recordings. Once the surgery was completed, an awake mix (Flumazenil (Anexate): 0.5 mg/kg, Atipamezol (Alzane): 2.5 mg/kg, Naloxone: 1.2 mg/kg) was injected subcutaneously before the animal was placed back in his cage with food and playing materials. Antibiotics (trimethoprime-sulfamethoxazole, Roche) and anti-inflammatory analgesics (Ibuprofen, Vifor, and Paracetamol, Bristol-Myers) were provided in the drinking water during the first 48 h. The animals were allowed to rest for 7 days and regain strength before performing kainate injections.

### Kainate and TTX injections

All injections were performed under isoflurane anesthesia. A small hole of about 0.3 mm was drilled above the site of injection using a nano inject (Drummond). Kainate or TTX was injected with a pulled glass capillary (Drummond glass capillaries # 3–000-203 G/X; tip diameter ∼15 µm) inserted through the brain. For kainate injections, a volume of 70 nl of kainate (Tocris Bioscience; 5 mM in NaCl 0.9%) was injected at a speed of 10 nl/s in the left dorsal hippocampus (Mediolateral 1.6 mm, Anteroposterior 1.8 mm, Depth 1.9 mm). A clear status epilepticus (SE) characterized by rearing and falling with forelimb clonus was observed in all animals. For TTX silencing, 0.15 µl of TTX (Sigma-Aldrich; 0.3 mM in NaCl 0.9%) was used to silence either the EF (AP −2.67 mm, ML 2.5 mm, depth 1.72 mm and 1.22 mm, angle 20°), the contralateral hippocampus (AP 2.54 mm, ML 2.10 mm, depth 2.0 for EEG; AP −2.67 mm, ML - 2.5 mm, depth 1.72 mm and 1.22 mm, angle 20° for intracortical experiments) or the ipsilateral frontal cortex (AP 1.98 mm, ML 1.3 mm, depth 0.5 mm). This procedure was shown to decrease efficiently neuronal activity (Sheybani et al., 2018) These regions were selected based on previous studies which showed the appearance of pathological epileptic activities in contralateral hippocampus and frontal cortex (Sheybani et al., 2018, 2019; Słowiński et al., 2019).

### Surface and intracortical EEG recordings

All animals were trained to remain in the head-fix position twice a day during 3-4 days before surface high-density EEG (HD-EEG) or bi-hippocampal intracortical recordings (iEEG). For all recording sessions, the animals were briefly anesthetized using light isoflurane anaesthesia to allow electrode positioning.Recordings were then performed once animals were completely awake. All recordings were acquired using a Digital Lynx SX (Neuralynx) and lasted for 30-60 minutes. HD-EEGs were recorded using 32-grid stainless steel electrodes covering the entire dorsal surface of the skull (see Fig. S1), as previously described (Mégevand et al., 2008; Sheybani et al., 2018), at a sampling rate of 4 KHz and a low pass of 2 KHz. Signals were re-referenced to the average reference offline for further analysis. Intra-cortical recordings were made with longitudinal 16 electrode probes (A16; Neuronexus) implanted in both hippocampi (AP, ML, depth of the lowermost channel, angles: 2.67, 2.5, 2.02, 20°; see Fig. S2), at a sampling rate of 16 KHz and a low pass of 8 KHz, referenced against an electrode located above the cerebellum. Data were analyzed with Cartool (D. Brunet, Center for Biomedical Imaging, University of Geneva, Switzerland) and custom-written scripts in MATLAB software (The Math Works).

### Detection and identification of epileptiform discharges

At the level of the EF, the classifications of epileptiform discharges varies across studies but a consensus emerges to consider the longer lasting (>5-10 sec) EF bursts with high spiking rates, that can be accompanied with behavioral immobility and muscular twitches (Bouilleret et al., 1999; Riban et al., 2002; Sheybani et al., 2018) as ictal-like patterns reminiscent of focal seizures, even if different terms have been used in the literature, such as “hippocampal paroxysmal discharges” (Riban et al., 2002; Sheybani et al., 2018), “high voltage sharp waves” (Twele et al., 2017; Welzel et al., 2020), “long epileptiform events” (Meier et al., 2007), “high-load bursts” (Heining et al., 2019) “electrographic seizures” (Klee et al., 2017) or simply focal/partial « seizures » (Bouilleret et al., 1999; Arabadzisz et al., 2005a). The shorter focal events are generally considered as interictal epileptiform discharges (IEDs) that we further classified here into classes of low or high paroxysmal discharges, using the consensual parameters of duration and intrinsic spiking frequencies. Secondary-generalized tonic-clonic electroclinical seizures are also present in the model (Bouilleret et al., 1999; Sheybani et al., 2018; Lisgaras and Scharfman, 2022), however at a lower rate of a few events per day. These secondary generalized seizures were more rarely detected in our recordings of 1h duration and therefore were not considered here.

To quantify epileptic events in a non-subjective manner, we applied a semi-automated procedure combining algorithm-based fast-ripples (FRs) detection and visually detected epileptiform discharges (blind to animal condition/stage).

#### Detection of focal epileptic activities

Our classification of epileptiform discharges generated in the EF matches most previous studies (Arabadzisz et al., 2005a; Krook-Magnuson et al., 2013; Twele et al., 2017; Sheybani et al., 2018; Heining et al., 2019) and is based on the spike load of the detected events. As described below, putative epileptiform discharges were first detected using fast-ripples (FRs) as a proxy and then further classified between interictal and ictal classes of epileptiform discharges based on their spike load, i.e. spike rate within the event, and their duration. FRs are not detected in control recordings or in saline-injected mice, except at low rates in the somatosensory areas (Baker et al., 2003; Sheybani et al., 2018). FRs occurring after kainate injection are thus considered as pathological. In both HD-EEG and iEEG recordings, we observed that overriding FRs almost systematically accompany epileptiform discharges, occurring during the spike component, and vice-versa (Weiss et al., 2016; Lévesque et al., 2018; Sheybani et al., 2018). The automated FRs detection algorithm (see below) therefore allows to standardize the detection of epileptiform discharges and eliminate any examiner bias toward the control or epileptic group, although it is possible that some epileptiform discharges not combined with FRs are discarded. We applied the same procedures for HD-EEG and iEEG epileptiform events detection to remain consistent. None of these spiking discharges were detected in recordings made before kainate injections (d-1). The absence of such epileptic activities in general, however not classified using the same methods than here, was previously demonstrated in saline injected mice (Twele et al., 2017; Sheybani et al., 2018).

#### Classification of focal epileptiform discharges

For HD-EEG, a marker is first positioned on the positive peak (in the raw data) of the spike waveform associated with each FR detected at electrodes above the EF (electrodes 17, 19, 20, 23, 24, see Fig. S1). For iEEG, the marker is positioned at the signal peak of the spike waveform in the DG (negative peak, see Fig. S2). As explained below, each FR is counted only once even when detected on several channels simultaneously. Inter-spike intervals (ISI) are then calculated as the peak-to-peak distances between spikes. These measures are then used to classify epileptic patterns in function of the spike rate (number of spikes/sec) and the event duration as follows (Fig. 1E). Discharges that were neither preceded nor followed by another spike for at least 1 sec were classified as isolated spikes (IS). Spiking discharges are defined as a single or a double spike waveform: if only two spikes are detected within one second, they belong to the same event and are classified as IS. Events with more than two successive spikes within a second were then further separated according to spike rates and event duration. Events that never display >5 spikes/second (for at least 1 second) are named <u>s</u>pike <u>t</u>rains (STs). The first detected spike that is not followed by another spike for at least one second is considered as the termination of the ST. These spike trains correspond to the high-voltage sharp waves (Riban et al., 2002), spikes trains (Maroso et al., 2011), short irregular spike trains (Arabadzisz et al., 2005b) or low- to medium-load spike bursts (Heining et al., 2019) described in previous studies. ST are thus characterized by a succession of spikes of intermediate density, not exceeding 5 spikes/sec and lasting around 2-5 sec. Trains of spikes that reach >5 spikes/second but do not exceed 10 seconds are classified as <u>s</u>hort <u>h</u>ippocampal <u>p</u>aroxysmal <u>d</u>ischarges (sHPDs). Trains of spikes that reach >5 spikes/second and last more than 10 seconds are classified as ictal <u>h</u>ippocampal <u>p</u>aroxysmal <u>d</u>ischarges (iHPDs). This type of electrographic paroxysmal event with high load of spikes and FRs are reminiscent of focal seizures, although concomitant epileptiform discharges can be observed in the contralateral hippocampus during some HPDs, and are generally classified as ictal events in the literature (Arabadzisz et al., 2005a; Krook-Magnuson et al., 2013; Twele et al., 2017; Sheybani et al., 2018).

As an indication, a total of 3755 IS, 710 STs, 134 iHPDs and 69 sHPDs were detected in 16 animals using 40-50 minutes iEGG in 16 animals. Generalized seizures that are accompanied by intense tonic-clonic spasms are also present in the IHK model (Lisgaras and Scharfman, 2022) but were more rarely detected in our recordings of 1h duration (only 3 events on iHPDs) and therefore were not separated here in a specific type.

#### Automated FRs detection algorithm

The same automated detector of FRs as previously published in (Sheybani et al., 2018; Słowiński et al., 2019), designed for both EEG and iEEG recordings, was used here to quantify focal and remote FRs. It recognizes occurrences of 4 consecutive oscillations with amplitude 3 times greater than the standard deviation of the 250 ms surrounding baseline. Before the screening, the data are filtered between 200 and 550 Hz (2^nd^ order Butterworth filter). To avoid identifying harmonics of lower frequency activities (e.g., high gamma) as FRs, the detector retained only those FRs that did not overlap with high-gamma activity. After that, all candidate events are visually validated in filtered and unfiltered data. A postprocessing analysis based on the phase lag between distinct events is performed to eliminate further FR caused by volume conduction at numerous contacts. If two events were identified at two different positions and if there was no phase lag between these two events, one could assume they resulted from volume conduction from the same generator. In such case only the event with the highest power was retained. The time between the first and last oscillations over the threshold is used to calculate the duration of FR. The intrinsic FR frequency is the frequency peak that occurred within 200-550 Hz of each occurrence. The sum of the absolute squares of each event is used to compute the power of FRs (normalized for the duration of the FR). Although FRs are often considered as markers of the EF, they can also be observed at distance of the EF. Note that remote FRs are more easily detected than remote spikes in HD-EEG. As described in (Sheybani et al., 2018, 2019), while remote spikes are not always easily visible in the raw signal, they are put into light when averaging signals locked on FRs marker, in the same way that event related potentials need numerous repetitions to be visible in surface EEG studies. Remote interictal FRs are therefore useful to quantify remote epileptiform discharges. For the FRs quantifications in specific regions of interests in the TTX experiments, we pooled the FRs detected at electrodes above the EF (electrodes 17, 19, 23, 24; see Fig. S1), the contralateral hippocampus (electrodes 3, 5, 8, 22) and the ipsilateral frontal cortex (electrodes 14, 28, 29, 30).

#### Detection of network IEDs

Network IEDs were identified visually and correspond to the previously described generalized spikes in (Sheybani et al., 2018); we decided to change their designation from generalized to network IEDs to better reflect the focal - and not generalized - nature of the model. These propagating IEDs invade a large-scale network and are characterized by two peaks of high amplitude lasting 30-40 milliseconds, separated by flattening of the EEG of approximately 250 ms. It has been previously described that the intracortical network IEDs onset is first driven by 2 predominant nodes in both the EF and the contralateral hippocampus before it invades frontal regions across both hemispheres (Sheybani et al., 2018). The morphology of these NETWORK IEDs were consistent between mice, and they were not observed either in the current control mice (d-1) or in previous studies using saline injected mice (Sheybani et al., 2018). We quantified NETWORK IEDs as separate events and did not include them in the focal IEDs described in the previous sections. Interictal spiking discharges of similar duration and invading both hippocampi were also observed in our iEEG recordings. Although these events are presumably reminiscent of network IEDs, these were not included in the current study. The statistical onset of the network IEDs were calculated using previously described methods (Sheybani et al., 2018). Briefly, the start of the network IED was set to the maximum value of the first peak of the global field power (GFP) (Brunet et al., 2011). Next, the statistical onset was calculated as the first time point that crossed the 5 SD threshold of the baseline, which was calculated from the 950 ms period before the network IED.

### Time-frequency Analysis

For the calculation of the background power, 50 epochs were selected for each recording such that they were 2 sec away from any detected epileptic events. Power was calculated as a function of frequency using ft_freqanalysis of the fieldtrip toolbox (Oostenveld et al., 2011) for each epoch and averaged across epochs. For iEEG data, we selected the channel with the maximum background power in the theta band (4-12 Hz), since this range dominates hippocampal activity. In the time-frequency plots (Fig. 1 Fig. S1), power was calculated using wavelet analyses (Morlet waves). The depicted results are z-scores with values < 2 set to 0. For the illustrative example of PSD from one mouse in Figure 6 A, 165 IS and 21 ST+HPD were included.

### Multi-unit activity (MUA) detection

MUA was detected in iEEG recordings based on a threshold method (Quiroga et al., 2004). We filtered the data between 600-6000 Hz to prevent overlapping with the frequency spectrum of FRs (200-550 Hz). The threshold for MUA detection was then specified as

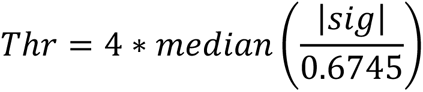

where sig denotes the entire signal filtered within MUA’s frequency band. To avoid detecting multiple action potentials on the same event, we optimized the detection by adding a “refractory” parameter that prevented from detecting another event within 1.8 ms (a compromise between the known duration of action potentials and several trials in our physiological recordings).

### Phase-locking and phase-synchronization

To decipher the role of bi-hippocampal network activities in the generation of ictal and interictal focal discharges, we performed phase locking and phase synchronization analysis in the periods preceding focal epileptiform discharges.

The statistical onset of an epileptic discharge was calculated based on the threshold detection method. The threshold was measured as

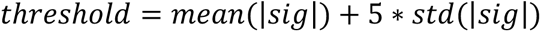

where sig is the baseline signal preceding the epileptiform discharge, from −1700 to −700 ms before the peak. The baseline offset is set at 300 ms before the epileptiform discharge peak to avoid any leakage effects from the spectral content of the epileptiform discharges. Over a time beginning 300 ms before the global field power (GFP) peak of the epileptiform discharges, our algorithm screened for onsets of LFP and ipsilateral and contralateral MUA. MUA activity was first convoluted to get a continuous signal using a Hanning window of 10 ms. To identify the statistical onset of LFP and MUA, we first evaluated the mean and standard deviation (SD) of either time series (LFP or convoluted MUA) over the baseline before peak of the ipsilateral epileptiform discharge as mentioned above. The statistical onset was defined as the timeframe when activity exceeded the threshold during 10 ms.

All phase-locking, phase synchronization and MUA analysis were performed in windows of 1 sec either preceding the statistical onsets of epileptiform discharges or in background periods. Data were first re-referenced to the cortical electrode, i.e., the uppermost electrode of the probe, positioned around the visual cortex L5, and filtered into 2 frequency bands (delta 0.5-4 Hz and theta 4-12 Hz). The filtering was done using the MATLAB filtfilt function, which is a non-causal filter, filtering the data in forward and backward directions and preventing the phase distortion of the filtered signal. The instantaneous amplitude and the phase are obtained by Hilbert-transform of the filtered signal using the MATLAB filter command Hilbert. The phase-locking of the epileptiform discharges to the slow-oscillations (SO) was obtained through intertrial coherence (ITC), which is defined as:

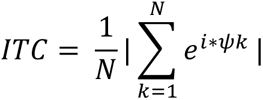

where N is the number of windows corresponding to specific epileptiform discharges or the background periods, k is the current window, and ψ is the phase of the signal. ITC was used across trials and within contacts to obtain phase-locking. Within each animal, ITC was calculated separately for each event and averaged across similar event type to obtain one ITC value for each animal and event type. We then averaged together ITC values of spike trains and hippocampal paroxysmal discharges (ST+HPD) within each animal for statistical comparisons. In both the EF and contralateral hippocampus probes, channels with the maximal theta power in the pyramidal region were selected for the statistical analysis of the phase-locking. For phase-synchronization, we used the maximal value of the ITC calculated between the channels of the pyramidal layer in the EF and CH. Statistical comparisons were made between the pre-epileptiform discharges periods and background periods, i.e. 1-second epochs spread across the entire recording, randomly selected such that these epochs are 2 seconds away from any epileptiform discharges and match the number of epileptic discharges in each recording session. The ratio of each measure (number of MUA, amplitude, phase-locking, and phase synchronization) was then calculated by dividing the value in the pre-epileptiform discharge period by the background value and one-sample Wilcoxon test was used to compare the ratio to median ratio set to 1. As per null hypothesis this ratio is equal to 1 if these is no difference between background and pre-IED period.

## References

1. D’Alessandro, M. et al. Epileptic seizure prediction using hybrid feature selection over multiple intracranial EEG electrode contacts: a report of four patients. IEEE Trans. Biomed. Eng. 50, 603–615 (2003).

2. Krook-Magnuson, E., Armstrong, C., Oijala, M. & Soltesz, I. On-demand optogenetic control of spontaneous seizures in temporal lobe epilepsy. Nature Communications 4, 1376 (2013).

3. Paz, J. T. et al. Closed-loop optogenetic control of thalamus as a tool for interrupting seizures after cortical injury. Nature Neuroscience 16, 64–70 (2013).

4. Sheybani, L. et al. Electrophysiological Evidence for the Development of a Self-Sustained Large-Scale Epileptic Network in the Kainate Mouse Model of Temporal Lobe Epilepsy. J. Neurosci. 38, 3776–3791 (2018).

5. Bragin, A., Wilson, C. L. & Engel, J. Chronic Epileptogenesis Requires Development of a Network of Pathologically Interconnected Neuron Clusters: A Hypothesis. Epilepsia 41, S144–S152 (2000).

6. Richardson, M. P. Large scale brain models of epilepsy: dynamics meets connectomics. J. Neurol. Neurosurg. Psychiatr. 83, 1238–1248 (2012).

7. Jehi, L. The Epileptogenic Zone: Concept and Definition. Epilepsy Curr 18, 12–16 (2018).

8. Weiss, S. A. et al. Graph theoretical measures of fast ripples support the epileptic network hypothesis. Brain Communications 4, (2022).

9. Bartolomei, F., Chauvel, P. & Wendling, F. Epileptogenicity of brain structures in human temporal lobe epilepsy: a quantified study from intracerebral EEG. Brain 131, 1818–1830 (2008).

10. Stead, M. et al. Microseizures and the spatiotemporal scales of human partial epilepsy. Brain 133, 2789–2797 (2010).

11. Bartolomei, F. et al. Defining epileptogenic networks: Contribution of SEEG and signal analysis. Epilepsia 58, 1131–1147 (2017).

12. Bouilleret, V. et al. Recurrent seizures and hippocampal sclerosis following intrahippocampal kainate injection in adult mice: electroencephalography, histopathology and synaptic reorganization similar to mesial temporal lobe epilepsy. Neuroscience 89, 717–729 (1999).

13. Riban, V. et al. Evolution of hippocampal epileptic activity during the development of hippocampal sclerosis in a mouse model of temporal lobe epilepsy. Neuroscience 112, 101–111 (2002).

14. Arabadzisz, D. et al. Epileptogenesis and chronic seizures in a mouse model of temporal lobe epilepsy are associated with distinct EEG patterns and selective neurochemical alterations in the contralateral hippocampus. Experimental Neurology 194, 76–90 (2005).

15. Sheybani, L., van Mierlo, P., Birot, G., Michel, C. M. & Quairiaux, C. Large-Scale 3–5 Hz Oscillation Constrains the Expression of Neocortical Fast Ripples in a Mouse Model of Mesial Temporal Lobe Epilepsy. eNeuro 6, (2019).

16. Słowiński, P. et al. Background EEG Connectivity Captures the Time-Course of Epileptogenesis in a Mouse Model of Epilepsy. eNeuro 6, ENEURO.0059–19.2019 (2019).

17. Mégevand, P. et al. A mouse model for studying large-scale neuronal networks using EEG mapping techniques. (2008) doi:10.1016/j.neuroimage.2008.05.016.

18. Weiss, S. A. et al. Ripples on spikes show increased phase-amplitude coupling in mesial temporal lobe epilepsy seizure-onset zones. Epilepsia 57, 1916–1930 (2016).

19. Lévesque, M., Salami, P., Shiri, Z. & Avoli, M. Interictal oscillations and focal epileptic disorders. European Journal of Neuroscience 48, 2915–2927 (2018).

20. Klee, R., Brandt, C., Töllner, K. & Löscher, W. Various modifications of the intrahippocampal kainate model of mesial temporal lobe epilepsy in rats fail to resolve the marked rat-to-mouse differences in type and frequency of spontaneous seizures in this model. Epilepsy Behav 68, 129–140 (2017).

21. Twele, F., Schidlitzki, A., Töllner, K. & Löscher, W. The intrahippocampal kainate mouse model of mesial temporal lobe epilepsy: Lack of electrographic seizure-like events in sham controls. Epilepsia Open 2, 180–187 (2017).

22. Heining, K. et al. Bursts with High and Low Load of Epileptiform Spikes Show Context-Dependent Correlations in Epileptic Mice. eNeuro 6, (2019).

23. Lisgaras, C. P. & Scharfman, H. E. Robust chronic convulsive seizures, high frequency oscillations, and human seizure onset patterns in an intrahippocampal kainic acid model in mice. Neurobiology of Disease 166, 105637 (2022).

24. Oostenveld, R., Fries, P., Maris, E. & Schoffelen, J.-M. FieldTrip: Open Source Software for Advanced Analysis of MEG, EEG, and Invasive Electrophysiological Data. Computational Intelligence and Neuroscience 2011, 1–9 (2011).

25. Jacobs, J. et al. High-frequency electroencephalographic oscillations correlate with outcome of epilepsy surgery. Annals of Neurology 67, 209–220 (2010).

26. Foffani, G., Uzcategui, Y. G., Gal, B. & Menendez de la Prida, L. Reduced Spike-Timing Reliability Correlates with the Emergence of Fast Ripples in the Rat Epileptic Hippocampus. Neuron 55, 930–941 (2007).

27. Avoli, M. et al. Specific imbalance of excitatory/inhibitory signaling establishes seizure onset pattern in temporal lobe epilepsy. J Neurophysiol 115, 3229–3237 (2016).

28. Sheybani, L. et al. Slow oscillations open susceptible time windows for epileptic discharges. Epilepsia 62, 2357–2371 (2021).

29. Alvarado-Rojas, C. et al. Slow modulations of high-frequency activity (40–140 Hz) discriminate preictal changes in human focal epilepsy. Sci Rep 4, 4545 (2015).

30. Haussler, U., Bielefeld, L., Froriep, U. P., Wolfart, J. & Haas, C. A. Septotemporal Position in the Hippocampal Formation Determines Epileptic and Neurogenic Activity in Temporal Lobe Epilepsy. Cerebral Cortex 22, 26–36 (2012).

31. Scharfman, H. E. & Myers, C. E. Hilar mossy cells of the dentate gyrus: a historical perspective. Front. Neural Circuits 6, (2013).

32. Bui, A. D. et al. Dentate gyrus mossy cells control spontaneous convulsive seizures and spatial memory. Science 359, 787–790 (2018).

33. Broggini, A. C. S., Esteves, I. M., Romcy-Pereira, R. N., Leite, J. P. & Leão, R. N. Pre-ictal increase in theta synchrony between the hippocampus and prefrontal cortex in a rat model of temporal lobe epilepsy. Experimental Neurology 279, 232–242 (2016).

34. Miller, J. W., Turner, G. M. & Gray, B. C. Anticonvulsant effects of the experimental induction of hippocampal theta activity. Epilepsy Research 18, 195–204 (1994).

35. Perucca, P., Dubeau, F. & Gotman, J. Intracranial electroencephalographic seizure-onset patterns: effect of underlying pathology. Brain 137, 183–196 (2014).

36. Lagarde, S. et al. Seizure-onset patterns in focal cortical dysplasia and neurodevelopmental tumors: Relationship with surgical prognosis and neuropathologic subtypes. Epilepsia 57, 1426–1435 (2016).

37. Sip, V., Scholly, J., Guye, M., Bartolomei, F. & Jirsa, V. Evidence for spreading seizure as a cause of theta-alpha activity electrographic pattern in stereo-EEG seizure recordings. PLOS Computational Biology 17, e1008731 (2021).

38. Aupy, J. et al. Cortico-striatal synchronization in human focal seizures. Brain 142, 1282–1295 (2019).

39. Chen, B. et al. A disinhibitory nigra-parafascicular pathway amplifies seizure in temporal lobe epilepsy. Nature Communications 11, 923 (2020).

40. Takeuchi, Y. et al. Closed-loop stimulation of the medial septum terminates epileptic seizures. Brain 144, 885–908 (2021).

41. Hristova, K. et al. Medial septal GABAergic neurons reduce seizure duration upon optogenetic closed-loop stimulation. Brain 144, 1576–1589 (2021).

42. Lopes, M. A., Junges, L., Woldman, W., Goodfellow, M. & Terry, J. R. The Role of Excitability and Network Structure in the Emergence of Focal and Generalized Seizures. Front. Neurol. 11, (2020).

43. Lévesque, M., Wang, S., Etter, G., Williams, S. & Avoli, M. Bilateral optogenetic activation of inhibitory cells favors ictogenesis. Neurobiology of Disease 171, 105794 (2022).

44. Mégevand, P. et al. Electric source imaging of interictal activity accurately localises the seizure onset zone. Journal of Neurology, Neurosurgery & Psychiatry 85, 38–43 (2014).

45. Kleen, J. K., Scott, R. C., Holmes, G. L. & Lenck-Santini, P. P. Hippocampal Interictal Spikes Disrupt Cognition in Rats. Ann Neurol 67, 250–257 (2010).

46. Kleen, J. K. et al. Hippocampal interictal epileptiform activity disrupts cognition in humans. Neurology 81, 18–24 (2013).

47. Avoli, M., Biagini, G. & de Curtis, M. Do Interictal Spikes Sustain Seizures and Epileptogenesis? Epilepsy Curr 6, 203–207 (2006).

48. Liou, J. et al. A model for focal seizure onset, propagation, evolution, and progression. eLife 9, e50927 (2020).

49. Smith, E. H. et al. Human interictal epileptiform discharges are bidirectional traveling waves echoing ictal discharges. eLife 11, e73541 (2022).

## Supporting material references

Arabadzisz D, Antal K, Parpan F, Emri Z, Fritschy J-M (2005a) Epileptogenesis and chronic seizures in a mouse model of temporal lobe epilepsy are associated with distinct EEG patterns and selective neurochemical alterations in the contralateral hippocampus. Experimental Neurology 194:76–90.

Arabadzisz D, Antal K, Parpan F, Emri Z, Fritschy J-MM, et al (2005b) Epileptogenesis and chronic seizures in a mouse model of temporal lobe epilepsy are associated with distinct EEG patterns and selective neurochemical alterations in the contralateral hippocampus. Experimental Neurology 194:76–90.

Baker SN, Gabriel C, Lemon RN, Stuart N. Baker * GC† and RNL‡ (2003) EEG oscillations at 600 Hz are macroscopic markers for cortical spike bursts. The Journal of Physiology 550:529–534.

Bouilleret V, Ridoux V, Depaulis A, Marescaux C, Nehlig A, Le Gal La Salle G (1999) Recurrent seizures and hippocampal sclerosis following intrahippocampal kainate injection in adult mice: electroencephalography, histopathology and synaptic reorganization similar to mesial temporal lobe epilepsy. Neuroscience 89:717–729.

Brunet D, Murray MM, Michel CM (2011) Spatiotemporal analysis of multichannel EEG: CARTOOL. Computational intelligence and neuroscience 2011:813870.

Bui AD, Nguyen TM, Limouse C, Kim HK, Szabo GG, Felong S, Maroso M, Soltesz I (2018) Dentate gyrus mossy cells control spontaneous convulsive seizures and spatial memory. Science 359:787–790.

Heining K, Kilias A, Janz P, Häussler U, Kumar A, Haas CA, Egert U (2019) Bursts with High and Low Load of Epileptiform Spikes Show Context-Dependent Correlations in Epileptic Mice. eNeuro 6 Available at: https://www.eneuro.org/content/6/5/ENEURO.0299-18.2019 [Accessed April 12, 2022].

Klee R, Brandt C, Töllner K, Löscher W (2017) Various modifications of the intrahippocampal kainate model of mesial temporal lobe epilepsy in rats fail to resolve the marked rat-to-mouse differences in type and frequency of spontaneous seizures in this model. Epilepsy Behav 68:129–140.

Krook-Magnuson E, Armstrong C, Oijala M, Soltesz I (2013) On-demand optogenetic control of spontaneous seizures in temporal lobe epilepsy. Nature Communications 4:1376.

Lévesque M, Salami P, Shiri Z, Avoli M (2018) Interictal oscillations and focal epileptic disorders. European Journal of Neuroscience 48:2915–2927.

Lisgaras CP, Scharfman HE (2022) Robust chronic convulsive seizures, high frequency oscillations, and human seizure onset patterns in an intrahippocampal kainic acid model in mice. Neurobiology of Disease 166:105637.

Maroso M, Balosso S, Ravizza T, Iori V, Wright CI, French J, Vezzani A (2011) Interleukin-1β Biosynthesis Inhibition Reduces Acute Seizures and Drug Resistant Chronic Epileptic Activity in Mice. Neurotherapeutics 8:304–315.

Mégevand P, Quairiaux C, Lascano AM, Kiss JZ, Michel CM, et (2008) A mouse model for studying large-scale neuronal networks using EEG mapping techniques.

Meier R, Häussler U, Aertsen A, Deransart C, Depaulis A, Egert U (2007) Short-term changes in bilateral hippocampal coherence precede epileptiform events. NeuroImage 38:138–149.

Oostenveld R, Fries P, Maris E, Schoffelen J-M (2011) FieldTrip: Open Source Software for Advanced Analysis of MEG, EEG, and Invasive Electrophysiological Data. Computational Intelligence and Neuroscience 2011:1–9.

Quiroga RQ, Nadasdy Z, Ben-Shaul Y (2004) Unsupervised Spike Detection and Sorting with Wavelets and Superparamagnetic Clustering. Neural Computation 16:1661–1687.

Riban V, Bouilleret V, Pham-Lê BT, Fritschy J-M, Marescaux C, Depaulis A (2002) Evolution of hippocampal epileptic activity during the development of hippocampal sclerosis in a mouse model of temporal lobe epilepsy. Neuroscience 112:101–111.

Sheybani L, Birot G, Contestabile A, Seeck M, Kiss JZ, Schaller K, Michel CM, Quairiaux C (2018) Electrophysiological Evidence for the Development of a Self-Sustained Large-Scale Epileptic Network in the Kainate Mouse Model of Temporal Lobe Epilepsy. J Neurosci 38:3776–3791.

Sheybani L, van Mierlo P, Birot G, Michel CM, Quairiaux C (2019) Large-Scale 3–5 Hz Oscillation Constrains the Expression of Neocortical Fast Ripples in a Mouse Model of Mesial Temporal Lobe Epilepsy. eNeuro 6 Available at: https://www.ncbi.nlm.nih.gov/pmc/articles/PMC6378326/ [Accessed March 20, 2019].

Słowiński P, Sheybani L, Michel CM, Richardson MP, Quairiaux C, Terry JR, Goodfellow M (2019) Background EEG Connectivity Captures the Time-Course of Epileptogenesis in a Mouse Model of Epilepsy. eNeuro 6:ENEURO.0059–19.2019.

Twele F, Schidlitzki A, Töllner K, Löscher W (2017) The intrahippocampal kainate mouse model of mesial temporal lobe epilepsy: Lack of electrographic seizure-like events in sham controls. Epilepsia Open 2:180–187.

Weiss SA, Orosz I, Salamon N, Moy S, Wei L, Van’t Klooster MA, Knight RT, Harper RM, Bragin A, Fried I, Engel J, Staba RJ (2016) Ripples on spikes show increased phase-amplitude coupling in mesial temporal lobe epilepsy seizure-onset zones. Epilepsia 57:1916–1930.

Welzel L, Schidlitzki A, Twele F, Anjum M, Löscher W (2020) A face-to-face comparison of the intra-amygdala and intrahippocampal kainate mouse models of mesial temporal lobe epilepsy and their utility for testing novel therapies. Epilepsia 61:157–170.

